# Circuit dysfunction in *SOD1-ALS* model first detected in sensory feedback prior to motor neuron degeneration is alleviated by BMP signaling

**DOI:** 10.1101/484337

**Authors:** Aaron Held, Paxton Major, Asli Sahin, Robert Reenan, Diane Lipscombe, Kristi A. Wharton

## Abstract

Amyotrophic lateral sclerosis (ALS) is a devastating neurodegenerative disease whose origin and underlying cellular defects are still not fully understood. While motor neuron degeneration is the signature feature of ALS, it is not yet clear if motor neurons, or other cells of the motor circuit, are the site of disease initiation. To better understand the contribution of multiple cell types in ALS, we made use of a *Drosophila Sod1^G85R^* knock-in model, in which all cells harbor the disease allele. End-stage *dSod1^G85R^* animals exhibit severe motor deficits with clear degeneration of motor neurons. Interestingly, earlier in *dSod1^G85R^* larvae motor function is also compromised, but their motor neurons exhibit only subtle morphological and electrophysiological changes, that are unlikely to cause the observed decrease in locomotion. We analyzed the intact motor circuit and identified a defect in sensory feedback that likely accounts for the altered motor activity of *dSod1^G85R^*. Furthermore, we found that the cell-autonomous activation of BMP signaling in proprioceptor sensory neurons that relay the contractile status of muscles back to the central nerve cord, is able to completely rescue early stage motor defects and partially rescue late stage motor function to extend lifespan. Identifying a defect in sensory feedback as a potential initiating event in ALS motor dysfunction, coupled with the ability of modified proprioceptors to alleviate such motor deficits, underscores the critical role that non-motor neurons play in disease progression and highlights their potential as a site to identify early-stage ALS biomarkers and for therapeutic intervention.

**Significance Statement:** At diagnosis, many cellular processes are already disrupted in the ALS patient. Identifying the initiating cellular events is critical for achieving an earlier diagnosis in order to slow or prevent disease progression. Our findings indicate that neurons relaying sensory information underlie early stage motor deficits in a *Drosophila* knock-in model of ALS that best replicates gene dosage in familial ALS (fALS). Importantly, studies on intact motor circuits revealed defects in sensory feedback before evidence of motor neuron degeneration. These findings strengthen our understanding of how neural circuit dysfunctions lead to neurodegeneration and coupled with our demonstration that activation of BMP signaling in proprioceptors alleviates both early and late motor dysfunction, underscores the importance of considering non-motor neurons as therapeutic targets.

## Introduction

Amyotrophic lateral sclerosis (ALS) is currently the most common motor neuron disease, at 1.9 cases per 100,000 people, with a projected global increase over the next 25 years (Chiò et al., 2013; Arthur et al., 2016). Most ALS patients experience a rapid decline in motor function due to the loss of motor neurons that ultimately results in respiratory failure and death within 3-5 years of diagnosis (Kiernan et al., 2011). Motor neurons of ALS patients exhibit changes in excitability, have cytoplasmic inclusions, and lose synaptic connectivity at the neuromuscular junction (NMJ) (Denys and Norris Jr., 1979; Tsujihata et al., 1984; Lowe et al., 1988; Kanai et al., 2006; Vucic et al., 2008; Iwai et al., 2016). Mutations in at least 50 genes have been linked to ALS (Taylor et al., 2016), many of which have broad expression domains encompassing many different cell types. The rapid and dramatic loss in the structural and functional integrity of motor neurons is typical of ALS, however, defects in glia, sensory neurons, cortical neurons, and skeletal muscle have all been noted in the literature (Tandan and Bradley, 1985; Theys et al., 1999; Zanette et al., 2002; Pugdahl et al., 2007; Taylor et al., 2016). Efforts to clarify the cellular origin of ALS-associated dysfunction will not only provide an understanding of the molecular and cellular basis of ALS progression, but will also inform new therapeutic strategies to treat, or even prevent motor neuron degeneration and loss.

In an effort to study the consequences of ALS-associated mutations in the context of the whole organism, where all affected cells are present and neuronal connections maintained, we made use of a *Drosophila Sod1^G85R^* knock-in model (Şahin et al., 2017). This *dSod1^G85R^* model harbors a mutation in the endogenous *dSod1* locus, synonymous with the human *SOD1^G85R^* mutation (Rosen et al., 1993), ensuring that its expression is physiologically relevant. *SOD1* is expressed ubiquitously, and *SOD1^G85R^* is compromised in its ability to eliminate the cellular build-up of superoxide while also causing a poorly understood toxic gain-of-function (Saccon et al., 2013; Şahin et al., 2017). Indicative of motor dysfunction, *dSod1^G85R^* adults fail to emerge from their pupal cases and subsequently die. Prior to this time, *dSod1^G85R^* 3^rd^ instar larvae exhibit reduced locomotion (Şahin et al., 2017). Larval crawling is a relatively simple behavior that requires a wave of contraction and relaxation of segmentally arranged muscles (Fox et al., 2006; Lahiri et al., 2011; Berni et al., 2012). A detailed knowledge of the larval motor circuit and the availability of genetic tools make *Drosophila* an ideal model to investigate the cellular and electrical basis of locomotor defects that arise from an ALS-causing mutation (Hughes and Thomas, 2007; Berni et al., 2012; Pulver et al., 2015).

Here, we report a detailed analysis of phenotypic defects associated with *dSod1^G85R^* at multiple time points in disease progression. Our data identify cells where defects first appear and a means to restore motor function in this ALS model. End-stage *dSod1^G85R^* animals, whose motor function is severely impaired, show a gross disruption in NMJ morphology that is accompanied by diminished neurotransmission. By contrast, the impaired locomotion of earlier-stage animals is not accompanied by a clear degeneration of the nerve-muscle synapse. Instead, we detect a disruption in non-motor neurons that mediate sensory feedback from the periphery to the interneuron network of the CNS. This defect acts to slow the overall locomotor pattern of *dSod1^G85R^* larvae. We also show that activation of Bone Morphogenetic Protein (BMP) signaling in non-motor neurons, as well as in motor neurons, can alleviate the abnormal motor function evident in both early- and end-stage *dSod1^G85R^* animals. Our findings implicate nonmotor neurons as causative in early-stage ALS-induced motor dysfunction, and as potential targets of therapeutic strategies aimed at preventing or slowing the ultimate degeneration of motor neurons.

## Materials and Methods

### Fly Stocks and Culture

The *w;*^+^*;dSod1^WTLoxP^* and *w;*^+^*;dSod1^G85R^/TM3SerGFP* lines (Şahin et al., 2017), *UAS-gbb^9.9^* (Khalsa et al., 1998), *UAS-SaxA* (Michael O’Connor), *OK6-Gal4* (BSC#64199, RRID:BDSC_64199), *OK371-Gal4* (P{GawB}VGlut^OK371^, BSC#26160, RRID:BDSC_26160), *MD-Gal4* (P{GawB}109(2)80, BSC#8769, RRID:BDSC_8769), *ChAT-Gal4* (P{ChAT-GAL4.7.4}, BSC#6798, RRID:BDSC_6798), *Repo-Gal4* (BSC#7415, RRID:BDSC_7415), *2-21-Gal4* (John Thomas), *UAS-mRFP* (Robert Reenan), *UAS-GFP.nls* (BSC#4775, RRID:BDSC_4775), and *BG57-Gal4* (33) were used to construct *w;*^+^*; dSod1^G85R^/TM6C, w; OK371-Gal4;dSod1^G85R^/TM6C, w;UAS-gbb^9.9^;dSod1^G85R^/TM6C, w;*^+^*;BG57-Gal4,dSod1^G85R^/TM6C, w;OK6-Gal4;dSod1^G85R^/TM6C, w;MD-Gal4;dSod1^G85R^/TM6C, w;*^+^*;2-21-Gal4;dSod1^G85R^/TM6C, w;UAS-mRFP;dSod1^G85R^/TM6C, w;OK371-Gal4;dSod1^WTLoxP^, w;^+^;2-21-Gal4;dSod1^WTLoxP^, w;UAS-mRFP;dSod1^WTLoxP^*, and *w;UAS-gbb^9.9^;dSod1^WTLoxP^*. Expression patterns for *Gal4* lines were found to be as previously reported (Budnik et al., 1996; Salvaterra and Kitamoto, 2001; Sepp et al., 2001; Mahr and Aberle, 2006; Hughes and Thomas, 2007; Sanyal, 2009). Flies were raised at 25°C on standard cornmeal/sugar/agar food. For all experiments, 2-3 males were crossed to 3 female virgins and brooded daily to control for progeny density. Both sexes of the experimental class were used to conduct experiments.

### Staging and Dissection

*Pharate NMJ:* Pharates (fully developed adults prior to eclosion) were staged based on the position of meconium along the anterior/posterior axis, with midway down the dorsal abdomen, corresponding to 9-10hrs before expected eclosion (Crossley, 1980). All pharates used for immunofluorescence or electrophysiology experiments were confirmed to be alive based on visible beating of the dorsal vessel. Pharate dissections essentially as described in Hebbar et al., 2006. Briefly, pharate was pulled from pupal case using forceps and placed in either 1xPBS (immunofluorescence experiments) or modified HL3.1 (electrophysiology). Pharate was pinned at its anterior and posterior ends dorsal side up and cut dorsally from posterior to anterior through the abdomen and thorax. Four additional pins were used to pin the pharate open, exposing the ventral muscles. Trachea, intestines, and reproductive organs were removed to expose the ventral muscles. Nerves connecting the VNC to body wall muscles were cut to allow clearer visualization of the body wall.

*Pharate Legs*: For leg extension assay, pharates were staged as for NMJ dissections and imaged using an Olympus SZX12 microscope with an OptixCam Summit D3K2-5 camera. Images were analyzed using ImageJ. Legs were dissected from pharates in PBS, mounted in 80% glycerol, and imaged using a Zeiss Axio Imager.M1 microscope. In the assignment of a leg integrity score, experimenters were blinded to the genotype and a score of nerve integrity (low (1) to high (5)) was assigned based on the continuity of the main nerve bundle and the number of collateral projections.

*3^rd^ instar larvae:* Wandering 3^rd^ instar larvae were identified as larvae >108hr AEL that had crawled out of the food and had not yet everted their anterior spiracles. Dissections were done either in PBS (immunofluorescence), modified HL3.1 (NMJ electrophysiology), or in a modified saline solution (fictive crawling and patching). Wandering 3^rd^ instar larvae were fileted as previously described (Jan and Jan, 1976; Zhang et al., 2010).

### Immunoflourescence

*Pharates and larvae*. Pharate or larval filets of the appropriate stage (see staging and dissection) were fixed for 15-20mins in 4% formaldehyde in 1xPBS (16% paraformaldehyde Electron Microscopy Sciences). After dissection and fixation, filets were blocked for 1 hour in 0.3% Normal Goat Serum (NGS) in 1xPBT (0.3% Triton-X100 in 1xPBS) at RT and then incubated overnight at 4°C with primary antibodies. Filets were then washed 2x5mins with 1xPBT and incubated for 1 hour in secondary antibody solution at RT. Filets were washed 2x5mins in 1xPBS, mounted in 80% glycerol with 0.5% N-propyl-gallate (Sigma-Aldrich) and imaged using a Zeiss LSM800 confocal microscope. The experimenter was blinded to genotype before counting boutons, and those counts represent the number of boutons per NMJ. The experimenter was also blinded while performing Sholl analysis and quantifying immunofluorescence. Primary antibodies and stains used in this study: mouse anti-Dlg (4F3, DSHB, RRID:AB_528203, at 1:200), rabbit anti-HRP-Cy3 (Jackson Immuno Research, RRID:AB_2340262, at 1:300), goat anti-HRP-647 (Jackson Immuno Research, RRID:AB_2338967, at 1:300), rabbit anti-pSmad3 (Abcam ab52903, RRID:AB_882596, at 1:300), rabbit anti-pSmad1 (Peter Ten Djike, at 1:1500), rabbit anti-RFP (Rockland, RRID:AB_2209751, at 1:500), mouse anti-BRP (nc82, DSHB, RRID:AB_2314866, at 1:50), chicken anti-GFP (Life Technologies A10262, RRID:AB_2534023, at 1:000), and phalloidin-488 (Life Technologies, RRID:AB_2315147, at 1:2500). Secondary antibodies: goat anti-mouse 647 (Life Technologies, RRID:AB_2535804, at 1:300), goat anti-chicken 488 (Life Technologies, RRID:AB_2534096, at 1:300), and goat anti-rabbit 568 (Life Technologies, RRID:AB_143157, at 1:300).

### Electrophysiology

*Adult NMJ*. Pharates were dissected from the pupal case (see staging and dissection) in modified HL3.1 (70mM NaCl, 5mM KCl, 4mM MgCl_2_, 10mM NaHCO_3_, 5mM Trehalose, 115mM Sucrose, 5mM HEPES, 1 mM CaCl_2_, pH 7.2). The four large ventral muscles (VM) in abdominal segment 2 (A2) were recorded from using 30MQ sharp electrodes filled with 3M KCl. After successfully entering one of the four large ventral muscles in A2, a 200pA current was injected for 250ms for 50 sweeps. The average sweep of the RC curves was used to determine the resistance and capacitance of the muscle. Only muscles with an input resistance >10MQ and a resting membrane potential < −40mV were used for analysis. For recordings of mini excitatory post-synaptic potentials (mEPSPs), negative current was injected to bring the membrane potential to approximately −55mV, and mEPSPs were recorded for 3 minutes. Recordings were made using a Multiclamp 700A, digitized with a Digidata 1550A, filtered at 10khz, and analyzed using custom Matlab scripts.

*Larval NMJ*. Wandering 3^rd^ instar larvae were fileted in modified HL3.1 (70mM NaCl, 5mM KCl, 10mM MgCl_2_, 10mM NaHCO_3_, 5mM Trehalose, 115mM Sucrose, 5mM HEPES, 0.5mM CaCl_2_, pH 7.2). Recordings at muscle 6 in segment A3 were done using 20-25MΩ sharp electrodes filled with 3M KCl. After successfully entering the muscle, a 1 nA current was injected for 150ms for 50 sweeps. The average sweep of the RC curves was used to determine the resistance and capacitance of the muscle. Only muscles with an input resistance >5MΩ and a resting membrane potential <-60mV were used for analysis. Evoked EPSPs were measured by suctioning a portion of the cut nerve into a suction electrode and passing a square 0.3ms pulse of current from a stimulation unit to evoke a compound action potential. A minimum of 10 ESPPs were averaged for each larva. mEPSPs were measured for 3 minutes in the absence of nerve stimulation. A second electrode was then inserted, muscle resistance was checked again, and the muscle was voltage clamped at −80mV. EPSCs were evoked by passing a 0.3ms pulse of current from a stimulation unit, and the gain was adjusted to reduce voltage error to <5mV. Recordings were made using an Axoclamp 2B, digitized with a Digidata 1322A, and analyzed using custom Matlab scripts. The experimenter was blinded during data acquisition and analysis.

*Muscle contraction*. Wandering 3^rd^ instar larvae were fileted in a modified saline solution (128mM NaCl, 2mM KCl, 35mM Sucrose, 4mM MgCl_2_, 5mM HEPES, 2mM CaCl_2_, pH 7.2). The severed main nerve bundle innervating an A3 hemisegment was suctioned into an electrode as in the NMJ experiments, and the nerve bundle was stimulated by applying square pulses of 0.3ms with the same current amplitude used in the larval NMJ experiments to evoke post-synaptic potentials. The nerve bundle was then stimulated for 200ms for three trials at 10hz, 25hz, 50hz, and 100hz. Body wall responses were observed using a Nikon FN1 microscope with an OptixCam Summit D3K2-5 camera at 105 frames/sec. The experimenter was blinded to genotype and stimulus frequency. The length of the relaxed muscle was compared to the length of the muscle at maximum contraction during the stimulation. The results of the three trials for each stimulus intensity were averaged.

*Fictive Crawling*. Larvae of the appropriate stage were dissected in a modified saline solution (128mM NaCl, 2mM KCl, 35mM Sucrose, 4mM MgCl_2_, 5mM HEPES, 3mM CaCl_2_, pH 7.2, based on Jan and Jan 1976) leaving the CNS intact. The majority of larvae began to fictive crawl, and larvae that did not crawl or respond to a light touch were discarded. A portion of one of the nerves innervating segments A3-A7 was tightly suctioned into an electrode filled with modified saline solution, and activity was recorded for 10 minutes. Bursts of nerve activity always correlated with peristaltic muscle activity (Cattaert et al., 2001; Fox et al., 2006). The paired fictive crawling recordings were done using the same modified saline solution, but with 1.5mM CaCl_2_ to reduce background activity when the CNS was isolated. After recording from intact preparations for 6 minutes, the CNS was removed from the body wall, a portion of the severed nerve attached to the VNC was suctioned into the electrode, and recording continued for another 6 minutes. Recordings were done using a Multiclamp 700A amplifier, AC filtered at 1hz, digitized with a Digidata 1550A, and analyzed using Clampfit 10.6. The frequency of bursting events was analyzed, but the amplitude of bursting events was not analyzed since it is highly dependent upon the resistance of the electrode-nerve junction.

*Motor neuron patching*. The central nervous system from wandering 3^rd^ instar larvae were dissected in modified saline solution (128mM NaCl, 2mM KCl, 35mM Sucrose, 4mM MgCl_2_, 5mM HEPES, 2mM CaCl_2_, pH 7.2) and glued (Vetbond) to a sylgard plate. After clearing the glial layer by applying 1% Protease XIV (Sigma) locally with a broken electrode using alternating positive and negative pressure, aCC motor neurons were identified by position and morphology (Choi et al., 2004; Marley and Baines, 2011) and patched with 3-5MΩ patch electrodes filled with internal solution (150mM KCH_3_SO_3_, 2mM MgCl_2_, 2mM EGTA, 5mM KCl, 20mM HEPES, pH 7.35). After light suction to enter whole-cell configuration, series resistance was compensated and current steps (-20pA to 160pA in steps of 10pA, 0.5s duration) were applied in triplicate. Sweeps that were influenced by large EPSPs were discounted from analysis. Cells were then voltage clamped at −70mV, and cell capacitance and series resistance were compensated. Spontaneous rhythmic currents (amplitude>300pA) were recorded for >1.5 minutes in gap free mode. Data was analyzed using custom Matlab scripts.

### Behavior

*Larval distance*. 2-5 larvae were placed on a 22cm diameter dish filled with 1% agarose. 90 second videos were taken using a Dinolite AM3111 camera and analyzed using custom Matlab scripts.

*Larval peristalsis*. Single larvae were placed on a 14cm dish filled with 1% agarose. 60 second videos were taken using a SZX12 stereomicroscope with an OpitxCam Summit D3K2-5 camera. The experimenter was blinded for the analysis, and videos were analyzed manually to determine the number of peristaltic contractions. The first 5 forward peristaltic motions were analyzed frame by frame to determine the crawling wave duration.

*Eclosion. dSod1^G85R^* mutants were identified by the absence of the balancer TM6C,SbTb. Pupal cases were counted 13 days after egg laying and identified as full or empty, indicating whether the adult successfully eclosed. Successful eclosion was verified by the observation of Sb+ flies.

### Experimental Design and Statistical Analysis

*Comparison of continuous data*. For comparisons of 2 genotypes, we used a student’s t-test, accounting for the variance and pairing of the data when necessary. Two-tailed t-tests where used to analyze most experiments. One-tailed tests were used when the hypothesis specified an effect direction. Graphs indicate samples (black dots), mean (vertical red line), and standard error (horizontal red lines). When a Shapiro-Wilk normality test indicated that the data was not normally distributed, we used a Wilcoxon rank sum test. When comparing >2 groups, a Holm-Bonferroni correction for multiple hypotheses was applied. ^*^, ^**^, and ^***^ signify p<0.05, 0.01, and 0.001, respectively.

*Comparison of categorical data*. For comparisons of categorical eclosion data, we used the Fisher’s exact test and a Holm-Bonferroni correction for multiple hypotheses when comparing >2 groups. Graphs represent samples (black dots) and median (red line). ^*^, ^**^, and ^***^ signify p<0.05, 0.01, and 0.001, respectively.

*Repeated measures data*. For comparisons of data sets with repeated measures, we assessed differences between genotypes using generalized linear mixed-effect models (GLME). Graphs represent individual samples (lighter lines), means (darker lines), and standard error (shaded area). For the analysis of action potential frequency, only the linear portion of the action potential response (0pA to 100pA) was used to model firing frequency. ^*^, ^**^, and ^***^ signify p<0.05, 0.01, and 0.001, respectively.

*Experimental design*. Sample size was predetermined by empirically estimating the variance and expected effect size of the parameter being measured. In all cases, N represents the number of individual animals, except in patch recording experiments where two different aCC motor neurons were recorded from the same VNC on three occasions.

*Table 1*. Experiments were conducted as described in the behavior subsection of the Material and Methods. All experimental values and statistical information can be found in Table 1 and the text of the results section. Values of N reflect individual animals. Fisher’s exact tests were used to compare genotypes and Holm-Bonferroni corrections were applied to account for multiple comparisons.

**Table 1.** *<u>gbb</u>* <u>expression partially rescues the</u> *<u>dSod1^G85R^</u>* <u>eclosion defect.</u> *dSod1^G85R^* adults fail to emerge from the pupal case and complete eclosion (Şahin et al., 2017). Expression of *gbb* using two different drivers that express in motor neurons (*OK371-Gal4* or *OK6-Gal4*) or muscles (*BG57-Gal4*) increases the percentage of *dSod1^G85R^* that eclose.† indicates experimental group is statistically different (p<0.05) from all *dSod1^G85R^* controls.

| Genotype | Eclosed | Total | Percent Eclosion ± 95% CI |
| --- | --- | --- | --- |
| *dSod1*<sup>WTLoxP</sup> | 194 | 204 | 95.1 ± 2.96 |
| *dSod1*<sup>G85R</sup> | 0 | 99 | 0.0 ± 0.0 |
| *UAS-gbb*<sup>9.9/+</sup>; *BG57-Gal4*, *dSod1*<sup>G85R</sup>/*dSod1*<sup>G85R</sup> | 27 | 307 | 8.79 ± 3.17 † |
| *BG57*, *dSod1*<sup>G85R</sup>/*dSod1*<sup>G85R</sup> | 0 | 95 | 0.0 ± 0.0 |
| *UAS-gbb*<sup>9.9/+</sup>; *dSod1*<sup>G85R</sup> | 0 | 201 | 0.0 ± 0.0 |
| *dSod1*<sup>WTLoxP</sup> | 397 | 400 | 99.3 ± 0.85 |
| *OK371-Gal4/UAS-gbb*<sup>9.9</sup>; *dSod1*<sup>WTLoxP</sup> | 428 | 453 | 94.5 ± 2.10 |
| *dSod1*<sup>G85R</sup> | 0 | 185 | 0.0 ± 0.0 |
| *OK371-Gal4/UAS-gbb*<sup>9.9</sup>; *dSod1*<sup>G85R</sup> | 10 | 173 | 5.78 ± 3.48 † |
| *OK371-Gal4/+*; *dSod1*<sup>G85R</sup> | 1 | 134 | 0.75 ± 1.46 |
| *UAS-gbb*<sup>9.9/+</sup>; *dSod1*<sup>G85R</sup> | 1 | 182 | 0.55 ± 1.07 |
| *dSod1*<sup>WTLoxP</sup> | 591 | 591 | 100 ± 0.0 |
| *dSod1*<sup>G85R</sup> | 0 | 163 | 0.0 ± 0.0 |
| *OK6-Gal4/UAS-gbb*<sup>9.9</sup>; *dSod1*<sup>G85R</sup> | 11 | 201 | 5.47 ± 3.14 † |
| *OK6-Gal4/+*; *dSod1*<sup>G85R</sup> | 0 | 205 | 0.0 ± 0.0 |
| *UAS-gbb*<sup>9.9/+</sup>; *dSod1*<sup>G85R</sup> | 3 | 211 | 1.40 ± 1.60 |

*Figure 1*. Experiments were conducted as described in the immunofluorescence and electrophysiology subsections of the methods. All experimental values and statistical information can be found in the text of the results section. Values of N reflect individual animals. Data were normally distributed, and t-tests accounting for the variance of the data were used to test differences between genotypes. A one-tailed t-test was used to compare *dSod1^G85R^* and *OK371-Gal4/UAS-gbb^9.9^; dSod1^G85R^* because the hypothesis being tested is whether *gbb* rescues *dSod1^G85R^* phenotypes. Holm-Bonferroni corrections were applied to account for multiple comparisons.

**Figure 1.**
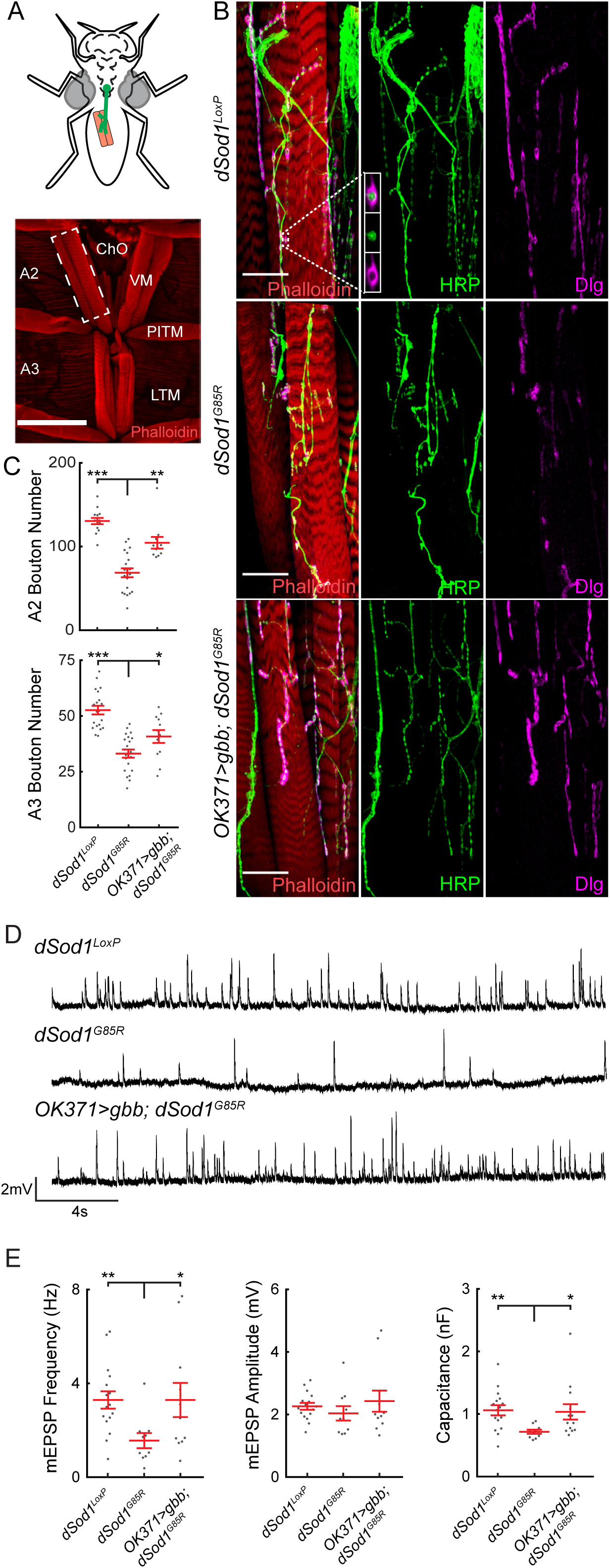
*dSod1^G85R^* mutants exhibit a reduction in bouton number, mEPSP frequency, and muscle capacitance in adult abdominal NMJs that can be rescued by the BMP ligand Gbb. **A**, Abdominal musculature of pharates is critical for eclosion. Ventral muscles in abdominal segments A2 and A3 shown (red, phalloidin) with ventral muscle (VM) NMJ in A2 hemisegment (white dashed box) shown in (**B**). **B**, High magnification of *dSod1^WTLoxP^, dSod1^G85R^*, and *OK371-Gal4/UAS-Gbb^9.9^; dSod1^G85R^* VM NMJ, presynaptic (green, anti-HRP), and post-synaptic membranes (magenta, anti-Dlg). Boutons are evident as presynaptic swellings along axonal branches. **C**, Bouton number is decreased in *dSod1^G85R^* mutants and increased in *OK371-Gal4/UAS-gbb^9.9^; dSod1^G85R^* VM NMJs. **D**, Representative traces of miniature excitatory postsynaptic potentials (mEPSPs) from *dSod1^WTLoxP^, dSod1^G85R^*, and *OK371-Gal4/UAS-Gbb^9.9^; dSod1^G85R^* VM. **E**, mEPSP frequency and muscle capacitance are decreased in *dSod1^G85R^*, mEPSP amplitude is not. Expression of *gbb* in glutamatergic neurons in *dSod1^G85R^* increases mEPSP frequency and muscle capacitance. Scale bars are 150μm (**A**) and 25μm (**B**).

*Figure 2*. Experiments were conducted as described in the Staging and Dissection subsections of the Material and Methods. All experimental values and statistical information can be found in the text of the results section. Values of N reflect individual animals. Leg extension data were normally distributed, and t-tests accounting for the variance of the data were used to test differences between genotypes. A one-tailed t-test was used to compare *dSod1^G85R^* and *OK371-Gal4/UAS-gbb^9.9^; dSod1^G85R^* because the hypothesis being tested is whether *gbb* rescues *dSod1^G85R^* leg extension. The nerve integrity score data is categorical, and was therefore analyzed using Fisher’s exact tests. Multiple comparisons were accounted for using Holm-Bonferroni corrections.

**Figure 2.**
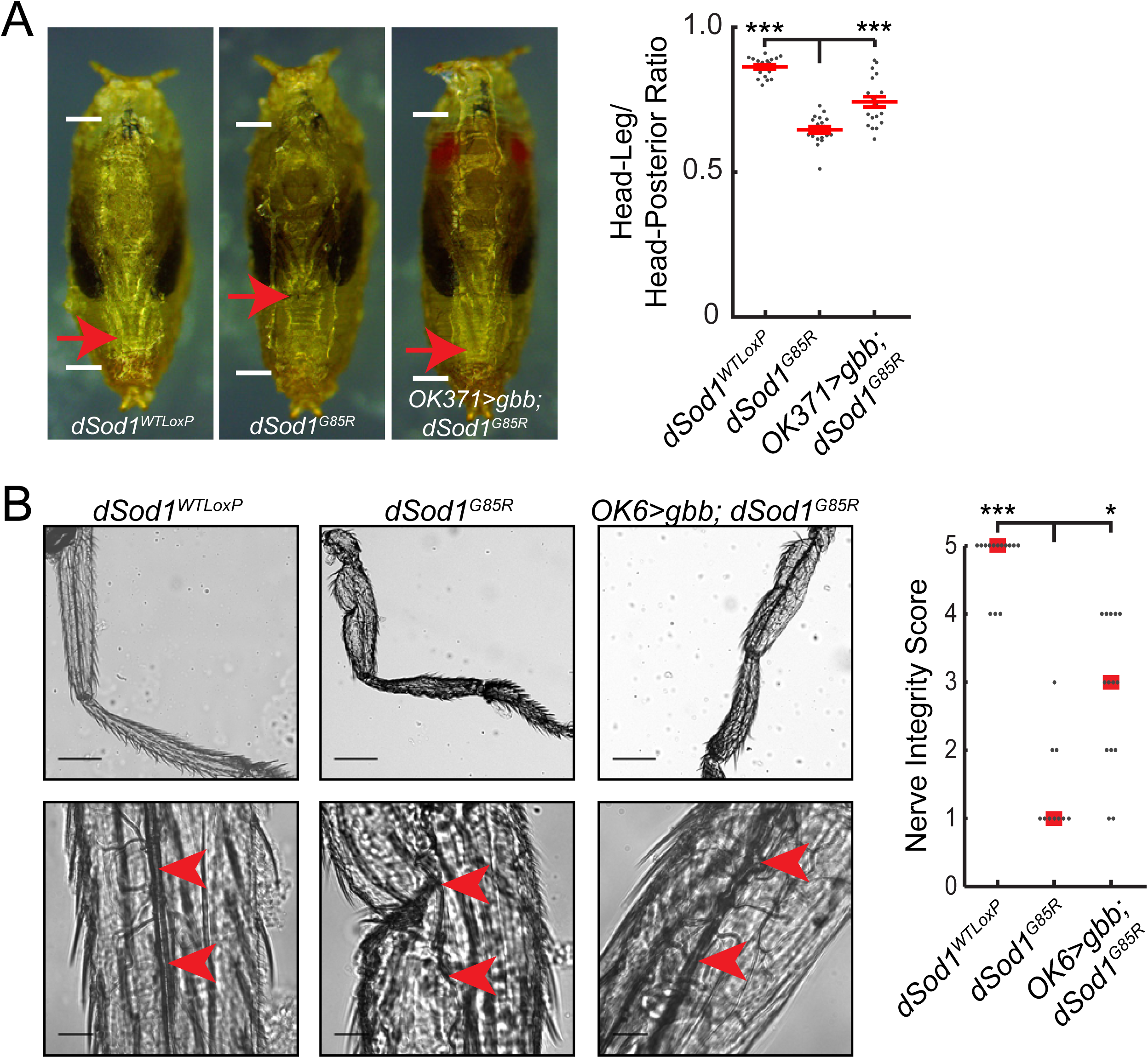
*gbb* expression rescues *dSod1^G85R^* leg extension and nerve integrity. **A**, Leg extension is compromised in *dSod1^G85R^* mutants. The failure of leg extension likely contributes to disruption in leg morphology (Şahin et al., 2017). The expression of *gbb* with *OK371-Gal4* partially rescues the *dSod1^G85R^* leg extension phenotype. White lines indicate pupal length, and red arrows indicate the tips of the metathoracic legs. **B**, *dSod1^G85R^* nerves (red arrowheads) in the metathoracic leg are severely disrupted and are rescued by expressing *gbb* with *OK6-Gal4*. Scale bars are 200μm in low mag images (**B**, top) and 30μm in high mag images (**B**, bottom).

*Figure 3*. Experiments were conducted as described in the Behavior and Electrophysiology subsections of the Material and Methods. All experimental values and statistical information can be found in the text of the results section. Values of N reflect individual animals. Data were normally distributed, and t-tests accounting for the variance of the data were used to test differences between genotypes. Holm-Bonferroni corrections were applied to account for multiple comparisons.

**Figure 3.**
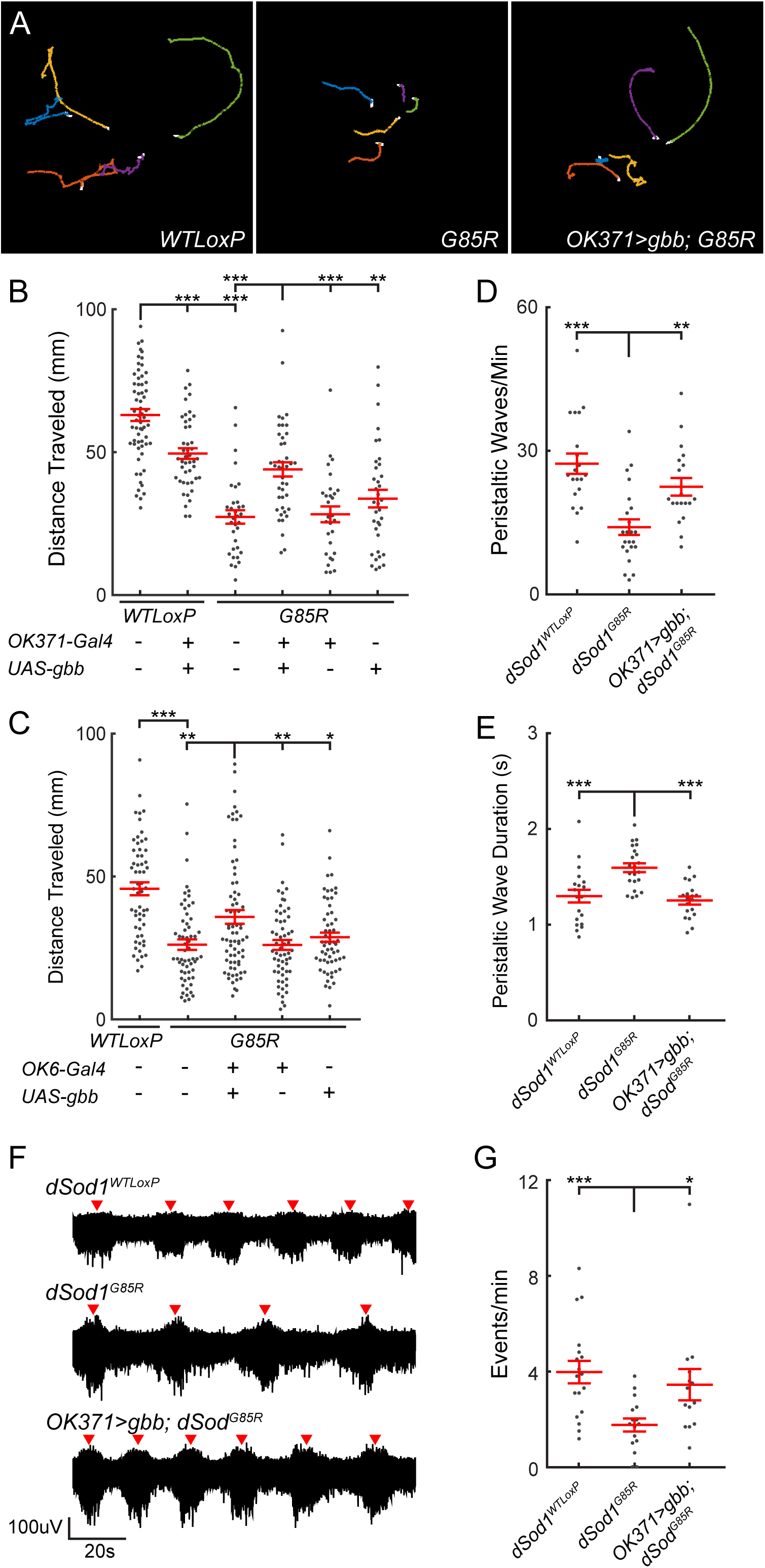
The *dSod1^G85R^* larval locomotion defect is improved by *gbb* expression. **A**, Tracks of wandering 3^rd^ instar larvae over 90s. **B**, Plot of distance traveled over 90s. Reduced locomotion of *dSod1^G85R^* larvae is increased by *OK371-Gal4* driven *gbb* (*OK371-Gal4/UAS-gbb^9.9^; dSod1^G85R^*) despite detrimental effects of *gbb* overexpression in wild type (*OK371-Gal4/UAS-gbb^9.9^; dSod1^WTLoxP^*). **C**, *gbb* expression using a second driver that also expresses in motor neurons, *OK6-Gal4*, also increases *dSod1^G85R^* larval locomotion. **D**, **E**, Quantification of number and duration of peristaltic waves of muscle contraction along the anterior/posterior axis of wandering 3^rd^ instar larvae. Waves, less frequent and longer in duration in *dSod1^G85R^*, are restored by *OK371-Gal4* driven *gbb. OK371-Gal4/UAS-gbb^9.9^; dSod1^G85R^* larvae have increased peristaltic wave frequency and decreased wave duration compared to *dSod1^G85R^*. **F**, **G**, Extracellular measurements of neural activity from segmental nerve in dissected larva with intact motor circuit. Bursts of neural activity (red arrowhead) are reduced in *dSod1^G85R^* reminiscent of the decrease in crawling wave frequency *in vivo. OK371-Gal4* driven *gbb* increases the frequency of events.

*Figure 4*. Experiments were conducted as described in the Immunofluorescence and Electrophysiology subsections of the Material and Methods. All experimental values and statistical information can be found in the text of the results section. Values of N reflect individual animals. Data were normally distributed in immunofluorescence measurements and NMJ recordings. T-tests accounting for the variance of the data were used to test differences between genotypes. Holm-Bonferroni corrections were applied to bouton counts to account for multiple comparisons. A generalized linear mixed-effect model was applied to measure genotype and stimulus effects on muscle contraction.

**Figure 4.**
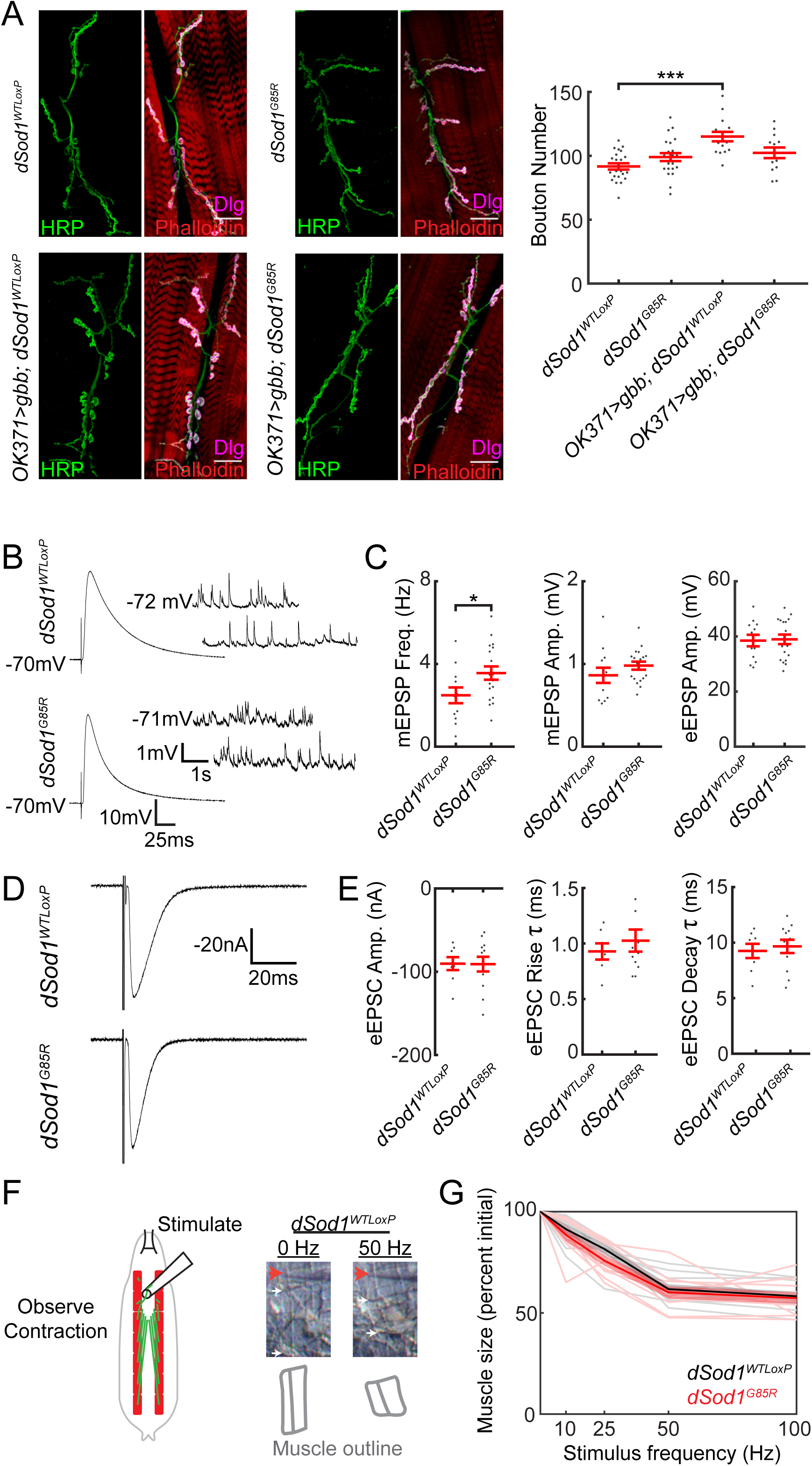
*dSod1^G85R^* larval NMJs exhibit only minor defects. **A**, Representative images of *dSod1^WTLoxP^, OK371-Gal4/UAS-gbb^9.9^; dSod1^WTLoxP^, dSod1^G85R^*, and *OK371-Gal4/UAS-gbb^9.9^; dSod1^G85R^* larval NMJs at muscle 6/7 segment A2. Muscles are stained with phalloidin (red), the presynaptic membrane is marked by anti-HRP (green), and the post-synaptic membrane is marked by anti-discs-large (Dlg) (magenta). Quantification of synaptic boutons, the presynaptic swellings that contain neurotransmitter, shows no significant difference between *dSod1^WTLoxP^* and *dSod1^G85R^*. The NMJ is expanded in *dSod1^WTLoxP^* but not in *dSod1^G85R^* when *gbb* is expressed in glutamatergic neurons under the control of *OK371-Gal4*. **B**, **C**, Electrophysiological recordings from *dSod1^WTLoxP^* and *dSod1^G85R^* show no significant change in eEPSP or mEPSP amplitude, but mEPSP frequency is increased in *dSod1^G85R^*. **D**, **E**, Voltage clamp recordings from *dSod1^WTLoxP^* and *dSod1^G85R^* show no significant changes in eEPSC amplitude, rise time, or decay time. The average of 10-15 tracings of eEPSP or eEPSC responses are shown. **F**, The A3 segmental nerve was stimulated for 200ms at 10hz, 25hz, 50hz, and 100hz and the extent of muscle contraction determined. The stimulus electrode (red arrow), segmental boundaries (white arrows), and outline of muscles 6/7 (gray) following 0Hz and 50Hz stimulation in *dSod1^WTLoxP^* are shown. **G**, Quantification of maximal muscle contraction (expressed as contracted muscle length/relaxed muscle length) in response to motor neuron stimulation (thick lines represent averages, thin lines represent individual muscles, and shaded areas represent standard error) shows no difference between *dSod1^WTLoxP^* (black, n=10) and *dSod1^G85R^* (red, n=10). Scale bars are 25 microns (**A**).

*Figure 5*. Experiments were conducted as described in the Electrophysiology subsection of the Material and Methods. All experimental values and statistical information can be found in the text of the results section. Values of N reflect patched cells. Most cells were taken from individual animals except on three occasions where two different aCC motor neurons were recorded from the same VNC. Data were normally distributed except for current density measurements. A Wilcoxon rank sum test was used to analyze current density data, generalized linear mixed-effect models were used to analyze repeated measure data, and t-tests were used to assess all other measurements.

**Figure 5.**
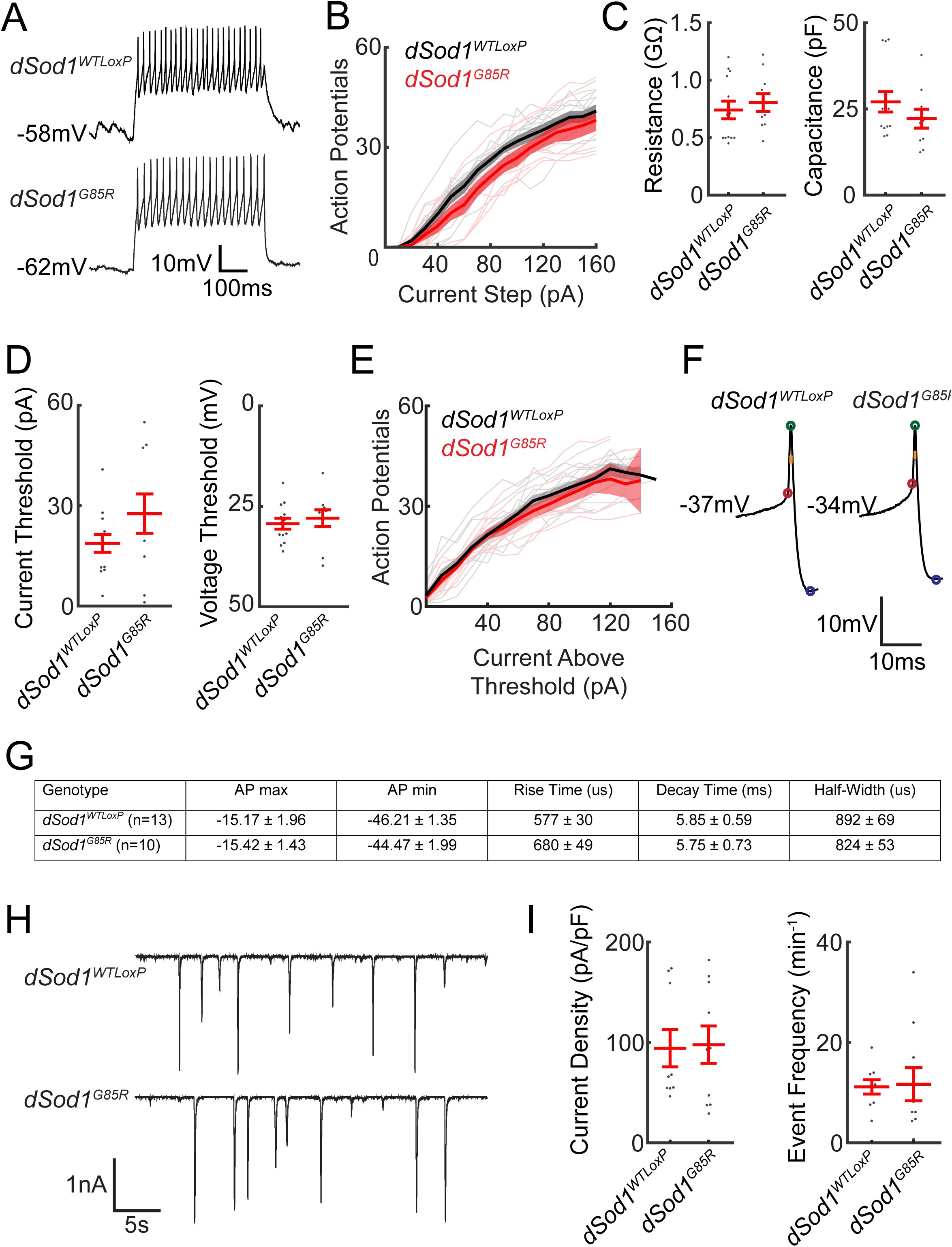
*dSod1^G85R^* larval motor neurons exhibit subtle defects. **A**, Action potential spikes in response to an 80pA current step are shown. **B**, Quantification of the action potential response to current steps from 0pA to +160pA for *dSod1^G85R^* (red) and *dSod1^WTLoxP^* (black) for individual aCC neurons (thin lines) are plotted with the average number of spikes per genotype (thick line) and standard error (shaded band) indicated. *dSod1^G85R^* motor neurons fire fewer action potentials than *dSod1^WTLoxP^* in response to current injection. **C**, Measurements of aCC input resistance and capacitance did not differ between *dSod1^G85R^* and *dSod1^WTLoxP^*. **D**, The current threshold in *dSod1^G85R^* showed a trend towards higher values, but a difference in voltage threshold between *dSod1^G85R^* and *dSod1^WTLoxP^* was not observed. **E**, Plot of action potential response normalized to current threshold shows no difference in action potential response between *dSod1^WTLoxP^* and *dSod1^G85R^*. **F**, **G**, Specific properties of individual action potential spikes near current threshold were analyzed and do not reveal significant differences between *dSod1^WTLoxP^* and *dSod1^G85R^* waveforms (voltage threshold, red; action potential peak, green; rise time, red to green; action potential minimum, blue; decay time, green to blue; half-width, orange). **H**, **I**, Voltage clamp recordings of spontaneous excitatory currents were not significantly different between *dSod1^G85R^* and *dSod1^WTLoxP^* regarding peak current density or event frequency.

*Figure 6*. Experiments were conducted as described in the Electrophysiology subsection of the Material and Methods. All experimental values and statistical information can be found in the text of the results section. Values of N reflect individual animals. Data were normally distributed, and t-tests accounting for variance were used to assess differences between genotypes. A one-tailed t-test was used to compare *dSod1^G85R^* and *OK371-Gal4/UAS-gbb^9.9^; dSod1^G85R^* because the hypothesis being tested is whether *gbb* rescues *dSod1^G85R^* event frequency. Paired t-tests were used in panel F because the changes between cut and intact preparations were measured within the same animals.

**Figure 6.**
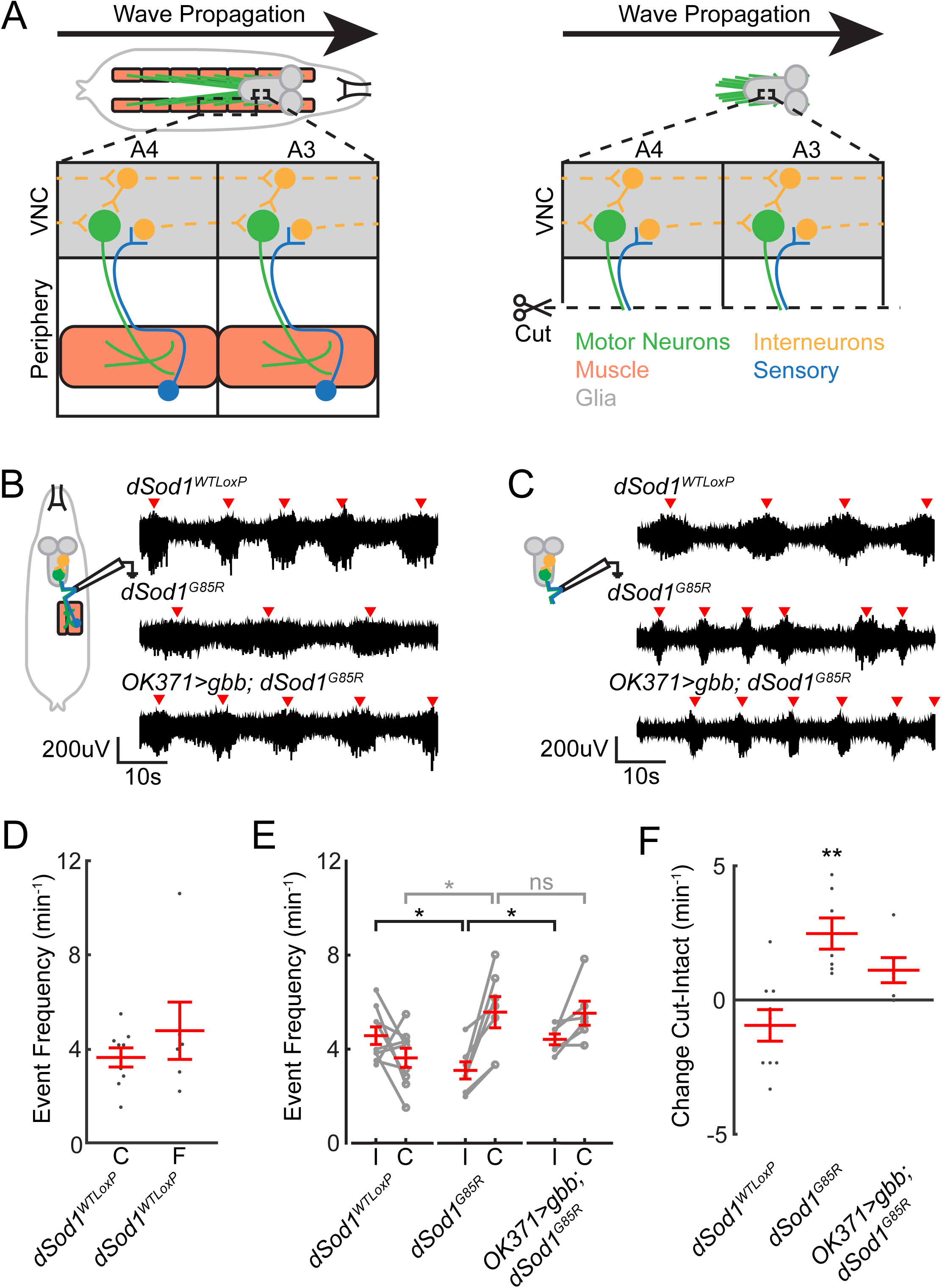
Peripheral feedback is disrupted in *dSod1^G85R^* larvae. **A**, Motor neurons (green) synapse onto muscles (orange). Sensory neurons (blue) detect stimuli, including muscle contraction, and feed back to the VNC. Interneurons (yellow) integrate sensory and central pattern generator (CPG) information within a segment and transfer signal to activate motor neurons in adjacent segment. Inputs to the interneuron network and CPG from the peripheral nervous system are removed by separating the brain/VNC from the body wall (right). **B**, Fictive crawling extracellular recordings from abdominal nerves in 3^rd^ instar larvae with an intact circuit, performed as in Figure 3. Red arrowheads mark bursts of neuronal activity. **C**, Extracellular recordings on abdominal nerves from isolated brain/VNC without input from peripheral feedback, after recordings were first performed with the circuit intact (**B**). **D**, Comparison of extracellular recordings from freshly dissected brains (“F”) with those that were performed on cut (“C”) nerves after first recording while the circuit was intact (ie., in **B**) showed no evidence of run-down on activity over the course of the experiment. **E**, Quantification of event frequency from paired intact “I” and cut “C” recordings. *dSod1^G85R^* exhibit a lower event frequency than *dSod1^WTLoxP^* when the circuit is intact. The lower event frequency of *dSod1^G85R^* is restored when *gbb* is expressed using *OK371-Gal4* (*OK371>gbb; dSod1^G85R^*). Neural activity in the isolated CNS, reflecting output from the CPG, is higher in *dSod1^G85R^* than *dSod1^WTLoxP^. OK371-Gal4* driven *gbb* does not affect this increase in CPG activity observed in *dSod1^G85R^*. **F**, Difference between each paired recording (Cut-Intact) indicates impact of peripheral feedback on circuit activity. Peripheral feedback does not significantly affect event frequency in *dSod1^WTLoxP^* or *OK371-Gal4/UAS-gbb^9.9^; dSod1^G85R^* but it slows the frequency of bursts in *dSod1^G85R^*. The slowing effect of peripheral feedback in *dSod1^G85R^* appears muted when *gbb* is expressed. The higher rate of bursting in *dSod1^G85R^* when feedback from the periphery is absent (**C**, **E**) is not altered by *OK371-Gal4* driven *gbb*, indicating that *gbb* expression does not overcome this defect.

*Figure 7*. Experiments were conducted as described in the Immunofluorescence subsection of the Material and Methods. All experimental values and statistical information can be found in the text of the results section. Values of N reflect individual animals. mRFP data was normally distributed and had unequal variance, so an unequal variance t-test was used to assess difference between genotypes. Sholl analysis data was analyzed using a generalized linear mixed effect model because it is a repeated measures data set.

**Figure 7.**
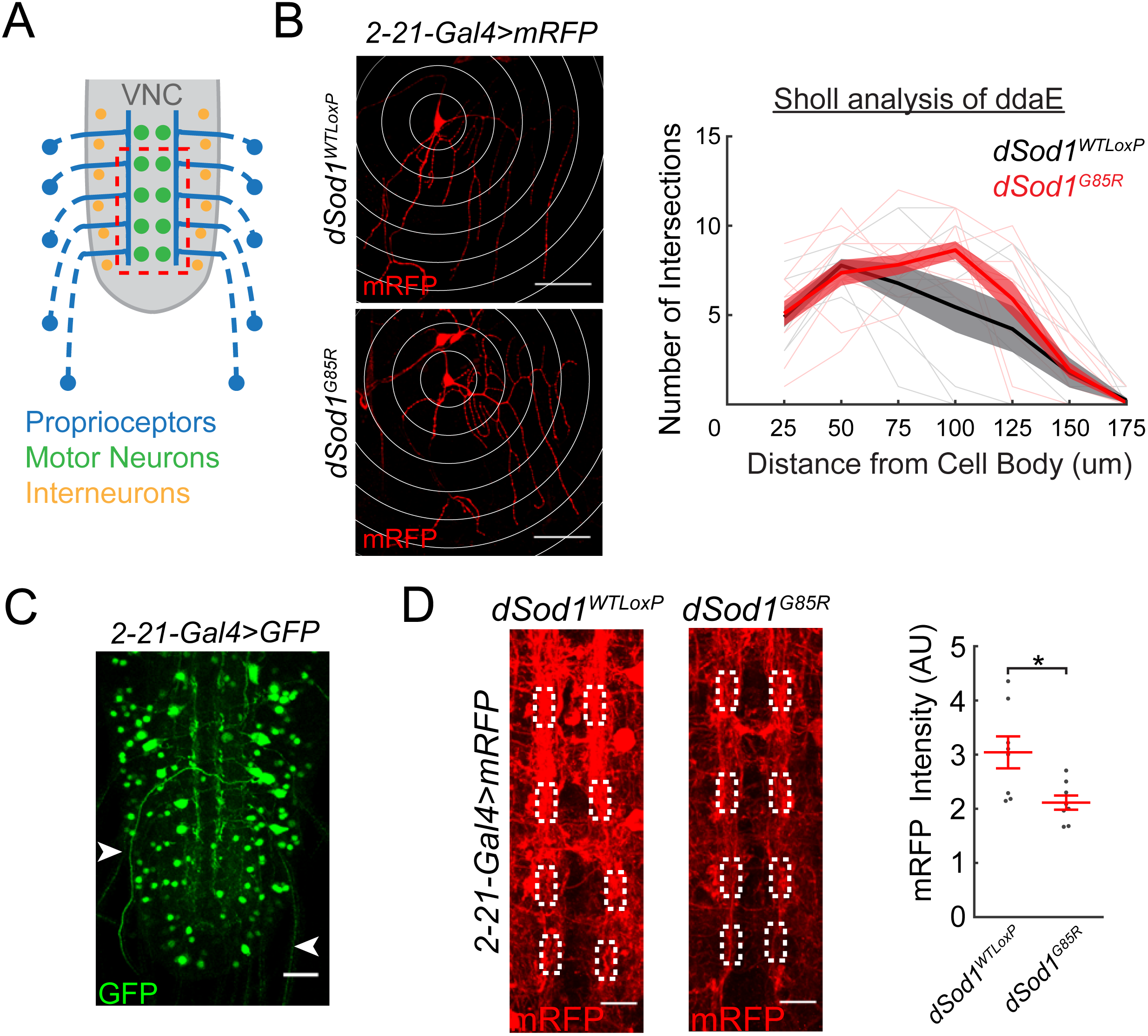
Neuronal processes are altered in *dSod1^G85R^* larval nerve cord. **A**, Schematic depicts axons of proprioceptors (blue) in the periphery projecting to the longitudinal tracts adjacent to the ventral nerve cord (gray) midline. **B**, The dendritic pattern of ddaE proprioceptors is marked by *2-21-Gal4>UAS-mRFP* and quantified using Sholl analysis (white rings, 25μm radius intervals) in both *dSod1^WTLoxP^* and *dSod1^G85R^* larvae. **C**, In addition to peripherally located proprioceptors, *2-21-Gal4* is also expressed in cells within the larval nerve cord, as indicated by *2-21-Gal4>UAS-GFP*. Axonal projections (white arrowheads) from multidendritic proprioceptor residing in the larval body wall are also visible. **D**, Axonal projections and other neuronal processes highlighted by *2-21-Gal4>UAS-mRFP* are apparent along the VNC midline (VNC midline shown corresponds to dashed red box in **A**) of *dSod1^WTLoxP^* and *dSod1^G85R^* late 3^rd^ instar larval VNCs. White boxes define the 5μm × 10μm regions of interest within the neuropil where proprioceptors project (Merritt and Whitington, 1995; Grueber et al., 2007), and the area of anti-mRFP intensity quantification. Scale bars are 50μm (**B**), 25μm (**C**), and 10μm (**D**).

*Figure 8*. Experiments were conducted as described in the Immunofluorescence subsection of the Material and Methods. All experimental values and statistical information can be found in the text of the results section. Values of N reflect individual animals. Data were normally distributed, and t-tests were used to assess differences between genotypes. A Holm-Bonferroni correction was applied to NMJ fluorescence measurements to account for multiple comparisons.

**Figure 8.**
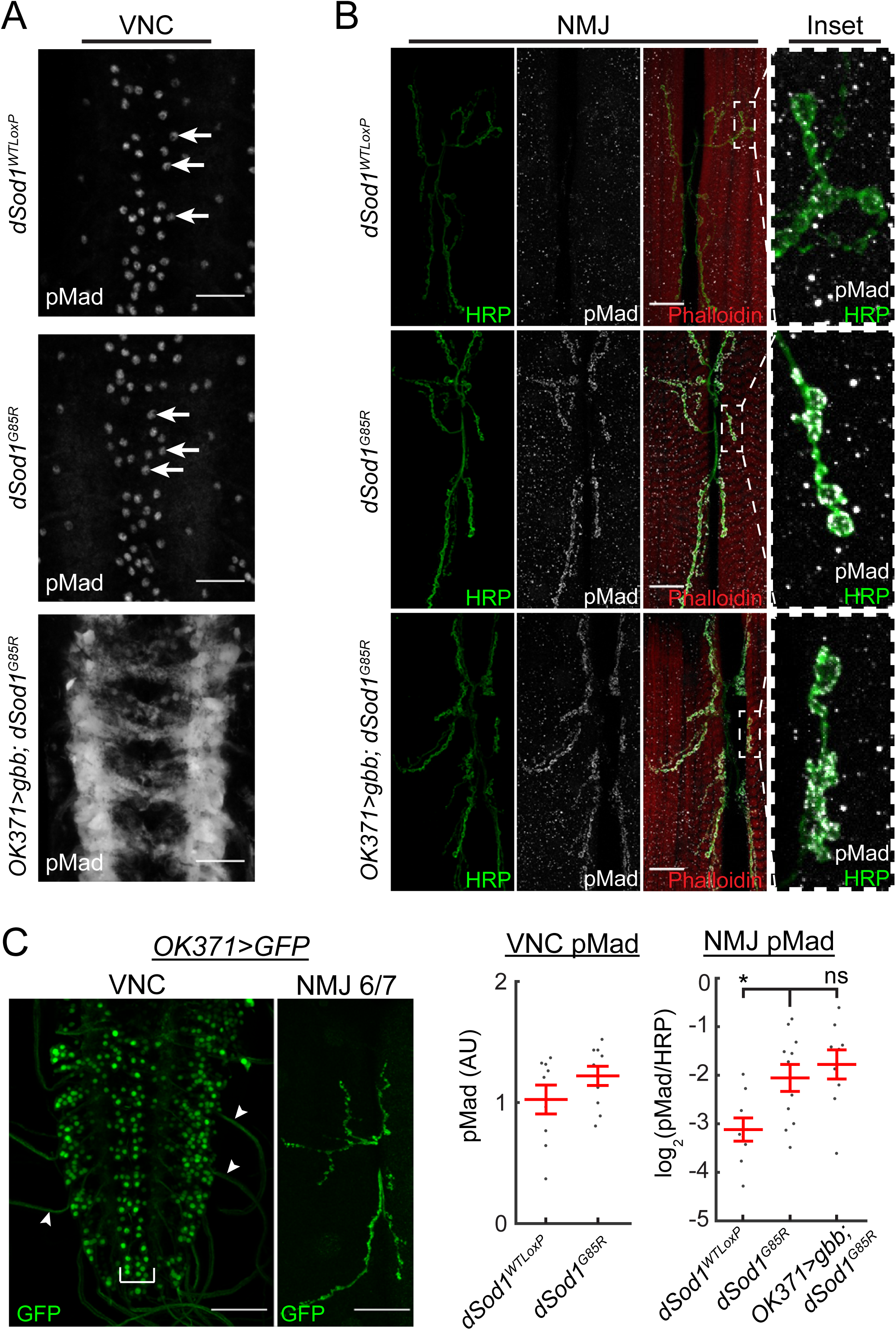
*gbb* expressed under the control of *OK371-Gal4* broadly induces pMad in VNC. **A**, pMad (gray) levels in motor neuron cell bodies (white arrows), as determined by anti-PS3 pMad antibody (Abcam ab52903, PS3), are unchanged in *dSod1^G85R^* compared to *dSod1^WTLoxP^*. The expression of *gbb* in *dSod1^G85R^* led to broadened and increased pMad staining in cells and processes along the nerve cord midline. pMad in the VNC recognized by a different antibody (anti-pSmad1, a gift from Peter tenDjike) gave a similar staining pattern (not shown). **B**, At the NMJ (6/7), pMad is significantly higher in *dSod1^G85R^* compared to *dSod1^WTLoxP^*. Overexpression of *gbb* in glutamatergic motor neurons does not lead to a further increase in synaptic pMad (*OK371-Gal4>UAS-gbb*); green, presynaptic anti-HRP; red, muscle, phalloidin; gray, anti-PS1). **C**, The expression pattern of *OK371-Gal4* is highlighted by *OK371-Gal4>UAS-GFP* in the medially located motor neurons (bracket) of the VNC, as well as in glutamatergic interneurons. Motor neuron innervation of muscles 6/7 is evident. Scale bars are 25μm (**A**,**B**) and 50μm (**C**).

*Figure 9*. Experiments were conducted as described in the Behavior subsection of the Material and Methods. All experimental values and statistical information can be found in the text of the results section. Values of N reflect individual animals. Data were normally distributed and t-tests, accounting for unequal variance where appropriate, were used to assess differences between genotypes. Multiple hypotheses were accounted for using Holm-Bonferroni corrections.

**Figure 9.**
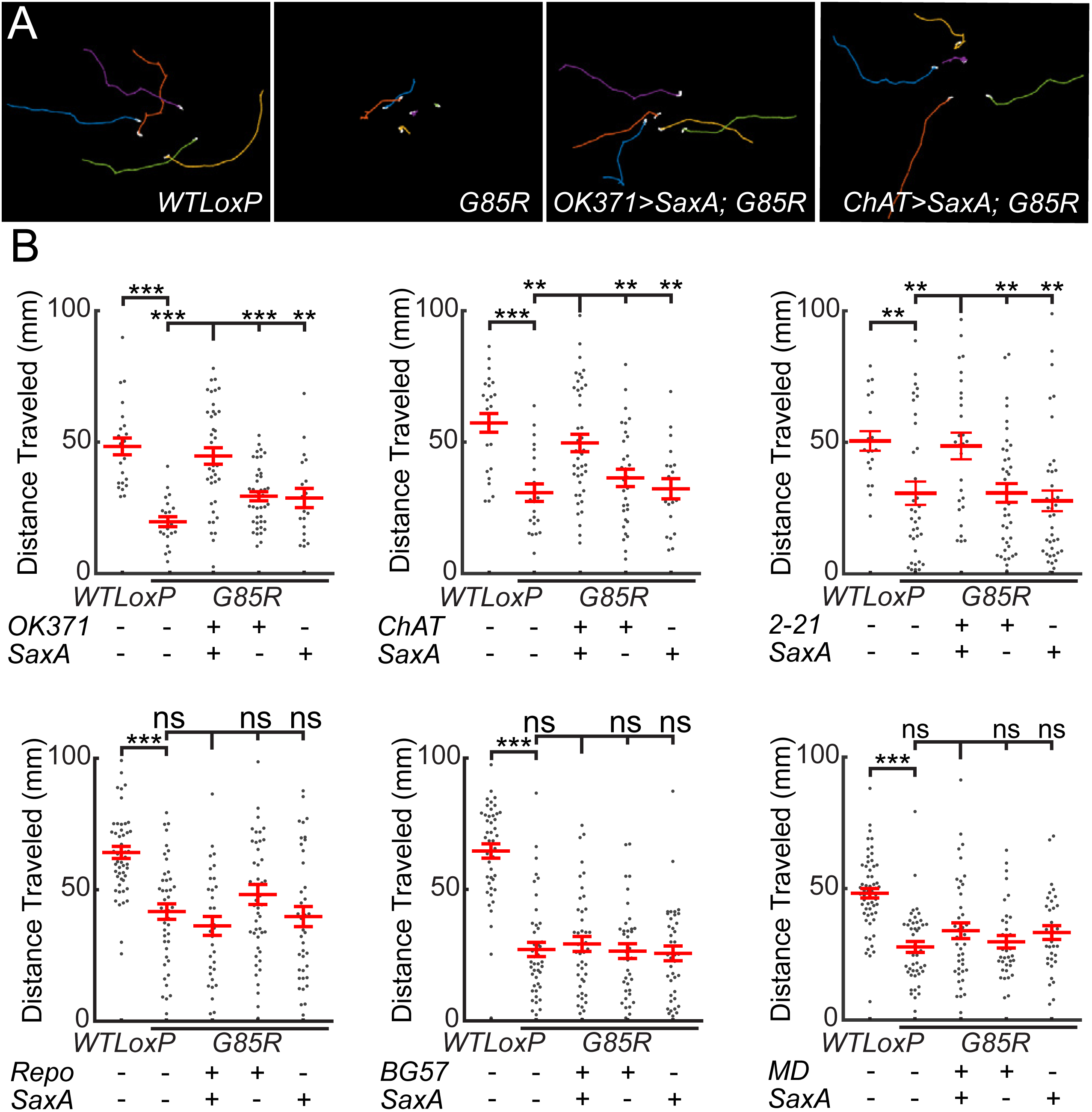
BMP signaling rescues *dSod1^G85R^* locomotion defect when activated in non-motor neurons. **A**, Traces of larvae crawling over 90 seconds. **B**, Cell autonomous activation expression of an activated BMP receptor, SaxA, in glutamatergic neurons (*OK371-Gal4*), cholinergic neurons (*ChAT-Gal4*), or proprioceptors (*2-21-Gal4*) rescues the larval locomotion defect of *dSod1^G85R^*. Expression in multidendritic sensory neurons (*MD-Gal4*), glial cells (*Repo-Gal4*) or muscle (*BG57-Gal4*) did not.

*Table 2*. Experiments were conducted as described in the Behavior subsection of the Material and Methods. All experimental values and statistical information can be found in Table 2 and the text of the results section. Values of N reflect individual animals. Fisher’s exact tests were used to compare genotypes and Holm-Bonferroni corrections were applied to account for multiple comparisons.

**Table 2.** <u>Cell-autonomous BMP signaling in cholinergic neurons rescues</u> *<u>dSod1^G85R^</u>* <u>eclosion.</u> Driving *UAS-SaxA* using *ChAT-Gal4* or *2-21-Gal4*, but not *OK371-Gal4, MD-Gal4, Repo-Gal4*, or *BG57-Gal4* increases the percent of *dSod1^G85R^* mutants that eclose.†, experimental group is statistically different (p<0.05) from all *dSod1^G85R^* controls.

| Genotype | Eclosed | Total | Percent Eclosion ± 95% CI |
| --- | --- | --- | --- |
| *dSod1<sup>WTLoxP</sup>* | 311 | 312 | 99.7 ± 0.61 |
| *dSod1<sup>G85R</sup>* | 0 | 106 | 0.0 ± 0.0 |
| *OK371-Gal4/+; UAS-SaxA, dSod1<sup>G85R</sup>/dSod1<sup>G85R</sup>* | 14 | 347 | 4.03 ± 2.07 |
| *OK371-Gal4/+; dSod1<sup>G85R</sup>* | 3 | 298 | 1.01 ± 1.13 |
| *UAS-SaxA, dSod1<sup>G85R</sup>/dSod1<sup>G85R</sup>* | 4 | 134 | 2.99 ± 2.88 |
| *dSod1<sup>WTLoxP</sup>* | 420 | 428 | 98.1 ± 1.28 |
| *dSod1<sup>G85R</sup>* | 0 | 104 | 0.0 ± 0.0 |
| *ChAT-Gal4/+; UAS-SaxA, dSod1<sup>G85R</sup>/dSod1<sup>G85R</sup>* | 41 | 401 | 10.2 ± 2.97 † |
| *ChAT-Gal4/+; dSod1<sup>G85R</sup>* | 2 | 213 | 0.94 ± 1.30 |
| *UAS-SaxA, dSod1<sup>G85R</sup>/dSod1<sup>G85R</sup>* | 3 | 138 | 2.17 ± 2.43 |
| *dSod1<sup>WTLoxP</sup>* | 509 | 513 | 99.2 ± 0.76 |
| *dSod1<sup>G85R</sup>* | 0 | 290 | 0.0 ± 0.0 |
| *2-21-Gal4, dSod1<sup>G85R</sup>/UAS-SaxA, dSod1<sup>G85R</sup>* | 96 | 270 | 35.6 ± 5.71 † |
| *2-21-Gal4, dSod1<sup>G85R</sup>/dSod1<sup>G85R</sup>* | 1 | 248 | 0.40 ± 0.79 |
| *UAS-SaxA, dSod1<sup>G85R</sup>/dSod1<sup>G85R</sup>* | 1 | 332 | 0.30 ± 0.59 |
| *dSod1<sup>WTLoxP</sup>* | 1321 | 1329 | 98.3 ± 1.16 |
| *dSod1<sup>G85R</sup>* | 0 | 555 | 0.0 ± 0.0 |
| *Repo-Gal4, dSod1<sup>G85R</sup>/UAS-SaxA, dSod1<sup>G85R</sup>* | 11 | 432 | 2.55 ± 1.49 |
| *Repo-Gal4, dSod1<sup>G85R</sup>/dSod1<sup>G85R</sup>* | 5 | 331 | 1.51 ± 1.31 |
| *UAS-SaxA, dSod1<sup>G85R</sup>/dSod1<sup>G85R</sup>* | 0 | 300 | 0.0 ± 0.0 |
| *dSod1<sup>WTLoxP</sup>* | 1063 | 1074 | 99.0 ± 0.60 |
| *dSod1<sup>G85R</sup>* | 0 | 462 | 0.0 ± 0.0 |
| *BG57-Gal4, dSod1<sup>G85R</sup>/UAS-SaxA, dSod1<sup>G85R</sup>* | 1 | 288 | 0.35 ± 0.68 |
| *BG57-Gal4, dSod1<sup>G85R</sup>/dSod1<sup>G85R</sup>* | 0 | 246 | 0.0 ± 0.0 |
| *UAS-SaxA, dSod1<sup>G85R</sup>/dSod1<sup>G85R</sup>* | 4 | 445 | 0.89 ± 0.87 |
| *dSod1<sup>WTLoxP</sup>* | 382 | 389 | 98.2 ± 1.32 |
| *dSod1<sup>G85R</sup>* | 0 | 217 | 0.0 ± 0.0 |
| *MD-Gal4/+; UAS-SaxA, dSod1<sup>G85R</sup>/dSod1<sup>G85R</sup>* | 2 | 171 | 1.17 ± 1.61 |
| *MD-Gal4/+; dSod1<sup>G85R</sup>* | 1 | 186 | 0.54 ± 1.05 |
| *UAS-SaxA, dSod1<sup>G85R</sup>/dSod1<sup>G85R</sup>* | 1 | 209 | 0.48 ± 0.94 |

All statistical comparisons were performed in MATLAB except for the multiple-category Fisher’s Exact test in Figure 2B, which was performed in R.

### Custom Scripts

The custom MATLAB scripts used for data analysis can be found at: https://github.com/ahheld/motor_circuit_dSod1ALS.

## Results

*End-stage dSod1^G85R^ mutants exhibit NMJ defects consistent with motor failure*

Homozygous *dSod1^G85R^* mutants die at the end of pupation, while wild type *dSod1^WTLoxP^* knock-in control animals emerge (eclose) from their pupal cases and live a normal adult lifespan (Şahin et al., 2017). The majority of *dSod1^G85R^* mutants die as fully formed adults that do not complete the eclosion process to emerge from the pupal case (pharates) (Table 1) (*dSod1^WTLoxP^* vs *dSod1^G85R^* p=3.1×10^−68^, Fisher’s Exact test with Holm-Bonferroni correction). Eclosion requires a series of abdominal muscle contractions, head thrusts and thoracic contractions to rupture the operculum and propel the adult fly from the pupal case (Kimura and Truman, 1990; McNabb et al., 1997). The failure of *dSod1^G85R^* mutants to eclose suggests a defect in muscle contractility.

We assessed the structural and functional integrity of abdominal muscle innervation, given their critical role in the eclosion process. Abdominal muscle fibers of *Drosophila* pharates are arranged in a stereotypical manner in each A2-A6 hemisegment (Figure 1A). We analyzed *dSod1^WTLoxP^* and *dSod1^G85R^* neuromuscular junctions (NMJs) at the ventral longitudinal muscles (VM) (Figure 1A) (Crossley, 1980; Kimura and Truman, 1990; Broadie and Bate, 1991). *dSod1^WTLoxP^* exhibit a characteristic arrangement of parallel axonal branches in the VM of A2 and A3 hemisegments, visualized by presynaptic (anti-HRP) and postsynaptic (anti-Dlg) markers (Sun and Salvaterra, 1995; Budnik et al., 1996) (Figure 1B). In contrast, the *dSod1^G85R^* A2 and A3 VM NMJs are disorganized, with kinked and discontinuous axonal projections. Furthermore, in *dSod1^G85R^* mutants the A2 and A3 NMJs contain fewer synaptic boutons, 53% and 63%, respectively, of *dSod1^WTLoxP^* controls (Figure 1C) (Segment A2: *dSod1^WTLoxP^* = 130.5 ± 3.8 boutons, n=14; *dSod1^G85R^* = 68.7 ± 5.3 boutons, n=20; *dSod1^WTLoxP^* vs *dSod1^G85R^:* p=2.2×10^−10^, unequal variance t-test with Holm-Bonferroni correction. Segment A3: *dSod1^WTLoxP^* = 52.6 ± 1.9 boutons, n=20; *dSod1^G85R^* = 33.1 ± 1.8 boutons, n=21; *dSod1^WTLoxP^* vs *dSod1^G85R^:* p=1.6×10^−8^, t-test with Holm-Bonferroni correction).

Consistent with the structural abnormalities of *dSod1^G85R^* NMJs, a substantial decrease in nerve muscle transmission in *dSod1^G85R^* pharates was evident compared to *dSod1^WTLoxP^* control animals (Figure 1D and E). The frequency of miniature excitatory postsynaptic potentials (mEPSPs) recorded from the VM in A2 hemisegments of *dSod1^G85R^* was 47% of that measured in *dSod1^WTLoxP^* (Figure 1E). mEPSP amplitudes measured in *dSod1^G85R^* and *dSod1^WTLoxP^* muscles were not distinguishable (Figure 1) <u>(mEPSP frequency</u>: *dSod1^WTLoxP^* = 3.29 ± 0.37s^−1^, n=16; *dSod1^G85R^* = 1.56 ± 0.32s^−1^, n=10; *dSod1^G85R^* vs *dSod1^WTLoxP^:* p=0.0071, t-test with Holm-Bonferroni correction; <u>mEPSP amplitude</u>: *dSod1^WTLoxP^* = 2.26 ± 0.11mV, n=16; *dSod1^G85R^* = 2.04 ± 0.23mV, n=10; *dSod1^WTLoxP^* vs *dSod1^G85R^:* p=0.67, t-test with Holm-Bonferroni correction). *dSod1^G85R^* VM muscle capacitance, 68% of *dSod1^WTLoxP^*, suggests that muscle surface area is reduced in *dSod1^G85R^* (*dSod1^WTLoxP^* = 1.06 ± 0.08nF, n=16; *dSod1^G85R^:* 0.73 ± 0.03nF, n=10; *dSod1^WTLoxP^* vs *dSod1^G85R^:* p=0.001, unequal variance t-test with Holm-Bonferroni correction). While we were unable to consistently evoke EPSPs in *dSod1^G85R^* preparations due to considerable motor neuron degeneration, the reduced frequency of spontaneous synaptic vesicle fusion events and the reduced number of boutons in *dSod1^G85R^* NMJs implicates presynaptic defects.

Abdominal and thoracic muscle contractions are also necessary to force hemolymph into everting pupal leg discs and extend newly formed legs (Mesce and Fahrbach, 2002). Consistent with a defect in abdominal and/or thoracic muscle contractility, *dSod1^G85R^* mutants fail to fully extend their legs (Figure 2A) (leg:body ratios: *dSod1^WTLoxP^:* 0.86 ± 0.01, n=19; *dSod1^G85R^:* 0.65 ± 0.01, n=20; p=2.0×10^−17^, unequal variance t-test with Holm-Bonferroni correction). The main nerve tracts of *dSod1^G85R^* legs are often discontinuous with a clear reduction in axonal arbors (Figure 2B and Şahin et al., 2017) (*dSod1^WTLoxP^* median score: 5, n=14; *dSod1^G85R^* median score: 1, n=10; *dSod1^WTLoxP^* vs *dSod1^G85R^* p=5.0×10^−5^, Fisher’s exact test with Holm-Bonferroni correction). These defects in both leg and abdominal muscle innervation, reduced NMJ and muscle size, and the associated failures in motor neuron dependent processes in *dSod1^G85R^*, are reminiscent of disruptions in muscle innervation, muscle wasting, and motor failure seen in other models of ALS and in human patients (Denys and Norris Jr., 1979; Tsujihata et al., 1984; Tandan and Bradley, 1985; Azzouz et al., 1997).

### BMP signaling enables dSod1^G85R^ eclosion

Members of the TGF-β/BMP family of signaling molecules act as neurotrophic factors to promote neuronal growth and survival (Jordan et al., 1997; Krieglstein et al., 2002; Lee-Hoeflich et al., 2004; Sun et al., 2007; Hocking et al., 2009). In *Drosophila*, the BMP7 orthologue, *gbb*, has previously been shown to promote NMJ growth and neurotransmission (McCabe et al., 2003; Goold and Davis, 2007; James et al., 2014). We tested the ability of Gbb, as a retrograde signal, to suppress motor defects observed in *dSod1^G85R^* mutants. Indeed, 8.8% of *dSod1^G85R^* flies successfully eclosed when Gbb was expressed in muscles (*UAS-gbb^9.9/+^; BG57-Gal4, dSod1^G85R^/dSod1^G85R^*) compared to 0% eclosion of *dSod1^G85R^* mutants (Table 1) (*dSod1^WTLoxP^* vs *dSod1^G85R^* p=3.1×10^−68^, *UAS-gbb^9.9/+^;BG57-Gal4,dSod1^G85R^/dSod1^G85R^* vs *dSod1^G85R^* p=1.4×10^−3^, *UAS-gbb^9.9/+^;BG57-Gal4,dSod1^G85R^/dSod1^G85R^* vs *BG57-Gal4,dSod1^G85R^/dSod1^G85R^* p=1.4×10^−3^ *UAS-gbb^9.9/+^;BG57-Gal4,dSod1^G85R^/dSod1^G85R^* vs *UAS-gbb^9.9/+^;dSod1^G85R^/dSod1^G85R^* p=4.6×10^−6^, Fisher’s Exact tests with Holm-Bonferroni correction). When *gbb* transcription is driven using two different *Gal4* lines that both express in motor neurons (OK371-Gal4 and OK6-Gal4; Mahr and Aberle, 2006; Sanyal, 2009) (*OK371-Gal4/UAS-gbb^9.9^; dSod1^G85R^* or *OK6-Gal4/UAS-gbb^9.9^; dSod1^G85R^*) a similar rescue was observed (Table 1) (*dSod1^WTLoxP^* vs *dSod1^G85R^* p=4.5×10^−151^, *OK371-Gal4/UAS-gbb^9.9^;dSod1^G85R^* vs *dSod1^G85R^* p=2.4×10^−3^, *OK371-Gal4/UAS-gbb^9.9^;dSod1^G85R^* vs *OK371-Gal4/+;dSod1^G85R^* p=0.026, *OK371-Gal4/UAS-gbb^9.9^;dSod1^G85R^* vs *UAS-gbb^9.9/+^;dSod 1^G85R^* p=0.014. *dSod1^WTLoxP^* vs *dSod1^G85R^* p=1.3×10^−169^, *OK6-Gal4/UAS-gbb^9.9^;dSod1^G85R^* vs *dSod1^G85R^* p=2.8×10^−3^, *OK6-Gal4/UAS-gbb^9.9^;dSod1^G85R^* vs *OK6-Gal4/+;dSod1^G85R^* p=1.1×10^−3^, *OK6-Gal4/UAS-gbb^9.9^;dSod1^G85R^* vs *UAS-gbb^9.9/+^;dSod1^G85R^* p=0.029, Fisher’s Exact tests with Holm-Bonferroni correction). In wild type animals, muscle-driven *gbb* results in an increase in body size while *OK371-Gal4* driven *gbb* does not (data not shown). In subsequent experiments aimed at understanding the impact of Gbb signaling on motor neuron function, we activated signaling by driving *gbb* expression with *OK371-Gal4*, not *BG57-Gal4*, to avoid the effects that a change in body size may introduce and confound our interpretation of results.

### Neuromuscular defects in end-stage dSod1^G85R^ suppressed by BMP signaling

We found that disruptions in the structural integrity of *dSod1^G85R^* VM NMJs, as well as their function were both rescued by *gbb* expression under the control of *OK371-Gal4* (*OK371>gbb; dSod1^G85R^*: *OK371 -Gal4/UAS-gbb^9.9^; dSod1^G85R^*) (Figure 1). Axonal projections and branching in *OK371>gbb; dSod1^G85R^* were restored to the characteristic parallel alignment of wild type VM NMJs (Figure 1B). An increase in bouton number was also observed at A2 and A3 VM NMJs (Figure 1C) with a corresponding increase in mEPSP frequency and muscle capacitance from recordings of *OK371>gbb; dSod1^G85R^* compared to *dSod1^G85R^* (Figure 1E) <u>(Segment A2 boutons</u>: *dSod1^G85R^* = 68.7 ± 5.3 boutons, n=20; *OK371-Gal4/UAS-gbb^9.9^; dSod1^G85R^* = 104.5 ± 7.0, n=11; *dSod1^G85R^* vs *OK371-Gal4/UAS-gbb^9.9^; dSod1^G85R^:* p=1.8×10-4, one-tailed t-test with Holm-Bonferroni correction. <u>Segment A3 boutons</u>: *dSod1^G85R^* = 33.1 ± 1.8 boutons, n=21; *OK371-Gal4/UAS-gbb^9.9^; dSod1^G85R^* = 40.8 ± 2.9, n=12; *dSod1^G85R^* vs *OK371-Gal4/UAS-gbb^9.9^; dSod1^G85R^:* p=0.012, one-tailed t-test with Holm-Bonferroni correction. <u>mEPSP frequency</u>: *dSod1^G85R^* = 1.56 ± 0.32s^−1^, n=10; *OK371-Gal4/UAS-gbb^9.9^; dSod1^G85R^* = 3.29 ± 0.73s^−1^, n=11; *dSod1^G85R^* vs *OK371-Gal4/UAS-gbb^9.9^; dSod1^G85R^:* p=0.024, one-sided unequal variance t-test with Holm-Bonferroni correction; <u>muscle capacitance</u>: *dSod1^G85R^:* 0.73 ± 0.03nF, n=10; *OK371-Gal4/UAS-gbb^9.9^; dSod1^G85R^:* 1.03 ± 0.12nF, n=13; *dSod1^G85R^* vs *OK371-Gal4/UAS-gbb^9.9^; dSod1^G85R^:* p=0.013, one-sided unequal variance t-test with Holm-Bonferroni correction).

In addition to the rescue of *dSod1^G85R^* abdominal NMJ phenotypes, *OK371>gbb; dSod1^G85R^* animals exhibited greater leg extension and improved leg nerve integrity compared to the mutants, consistent with a *gbb*-mediated improvement in abdominal muscle contractions and reduced leg motor neuron degeneration (Figure 2) <u>(leg extension</u>: *dSod1^G85R^:* 0.65 ± 0.01, n=20; *OK371-Gal4/UAS-gbb^9.9^; dSod1^G85R^* head:leg ratio = 0.74 ± 0.02, n=20; p=3.9×10^−5^ unequal variance one-tailed t-test with Holm-Bonferroni correction; <u>nerve integrity</u>: *dSod1^G85R^* median score: 1, n=10; *OK6-Gal4/UAS-gbb^9.9^; dSod1^G85R^* median score: 3, n=14; *dSod1^G85R^* vs *OK6-Gal4/UAS-gbb^9.9^; dSod1^G85R^* p=0.026, Fisher’s exact test with Holm-Bonferroni correction).

Thus, expression of the BMP ligand, Gbb, in a subset of neurons that includes motor neurons (*OK371>gbb; dSod1^G85R^*) alleviates defects in axonal branching, bouton number, mEPSP frequency, and muscle membrane capacitance at the *dSod1^G85R^* VM NMJs, as well as defects in leg extension and leg nerve integrity. However, despite the nearly complete restoration of neurotransmission across *OK371>gbb; dSod1^G85R^* abdominal NMJs, relatively few individuals fully eclose (5.8%), indicating that while the level of BMP signaling achieved by *OK371-Gal4* driven *gbb* expression is sufficient to greatly improve *dSod1^G85R^* NMJ function, other abnormalities that influence successful eclosion must not be fully restored.

### *Reduced locomotion of* dSod1^G85R^ *larvae is rescued by neuronal-driven* gbb

Like pharates, wandering 3^rd^ instar *dSod1^G85R^* larvae exhibit compromised muscle contraction. They crawl approximately half the distance of wild type (*dSod1^WTLoxP^*) animals during a 90 second period (Figure 3A-C) (*dSod1^WTLoxP^* = 63.0 ± 2.1mm, n=63; *dSod1^G85R^* = 27.3 ± 2.4mm, n=34, p=2.4×10^−17^, t-test with Holm-Bonferroni correction, and Sahin et al. 2017), with fewer posterior to anterior peristaltic waves of muscle contraction, each longer in duration, compared to *dSod1^WTLoxP^*(Figure 3D, E) <u>(Frequency</u>: *dSod1^WTLoxP^* = 27.3 waves/min, n=20; *dSod1^G85R^* = 14.0 waves/min, n=23; p=2.1×10^−5^, t-test with Holm-Bonferroni correction. <u>Duration</u>: *dSod1^WTLoxP^* = 1.30 ± 0.07s, n=20; *dSod1^G85R^:* 1.59 ± 0.05s, n=22; p=6.4×10^−4^, t-test with Holm-Bonferroni correction). When *gbb* is expressed under the control of *OK371-Gal4* or *OK6-Gal4, dSod1^G85R^* larvae travel farther (Figure 3A-C) (*dSod1^WTLoxP^* = 63.0 ± 2.1mm, n=63; *dSod1^G85R^* = 27.3 ± 2.4 mm, n=34; *OK371-Gal4/UAS-gbb^9.9^; dSod1^G85R^* = 44.0 ± 2.5mm, n=42; *OK371-Gal4/+^9^; dSod1^G85R^* = 28.2 ± 2.8mm, n=29; *UAS-gbb^9.9/+^; dSod1^G85R^* = 33.7 ± 3.1mm, n=36; *dSod1^WTLoxP^* vs *dSod1^G85R^* p=2.4×10^−17^; *OK371-Gal4/UAS-gbb^9.9^; dSod1^G85R^* vs *dSod1^G85R^* p=2.7×10^−5^; *OK371 -Gal4/UAS-gbb^9.9^; dSod1^G85R^* vs *OK371-Gal4/+^9^; dSod1^G85R^* p=1.7×10^−4^; *OK371 -Gal4/UAS-gbb^9.9^; dSod1^G85R^* vs *UAS-gbb^9.9^/+; dSod1^G85R^* p=0.011; t-tests with Holm-Bonferroni corrections. *dSod1^WTLoxP^* = 45.8 ± 2.3mm, n=58; *dSod1^G85R^* = 26.2 ± 1.8mm, n=60; *OK6-Gal4/UAS-gbb^9.9^; dSod1^G85R^* = 35.9 ± 2.4mm, n=73; *OK6-Gal4/+^9^; dSod1^G85R^* = 26.1 ± 1.7mm, n=59; *UAS-gbb^9.9^/+; dSod1^G85R^* = 28.8 ± 1.6mm, n=59; *dSod1^WTLoxP^*vs *dSod1^G85R^* p=2.8×10^−9^; *OK6-Gal4/UAS-gbb^9.9^; dSod1^G85R^* vs *dSod1^G85R^* p=9.3×10^−3^; *OK6-Gal4/UAS-gbb^9.9^; dSod1^G85R^* vs *OK6-Gal4/+^9^; dSod1^G85R^* p=0.017; *OK6-Gal4/UAS-gbb^9.9^; dSod1^G85R^* vs *UAS-gbb^9.9^/+; dSod1^G85R^* p=4.9×10^−3^; t-tests with Holm-Bonferroni corrections). An increase in the frequency of peristaltic waves relative to *dSod1^G85R^* mutants and a corresponding decrease in the duration of each wave was observed when *gbb* was expressed in glutamatergic neurons under the control of *OK371-Gal4* (Figure 3D and E) <u>(Frequency</u>: *dSod1^G85R^* = 14.0 waves/min, n=23; *OK371-Gal4/UAS-gbb^9.9^; dSod1^G85R^* = 22.47 ± 1.84 waves/min, n=19; p=6.9×10^−4^, onesided t-test with Holm-Bonferroni correction. <u>Duration</u>: *dSod1^G85R^:* 1.59 ± 0.05s, n=22; *OK371-Gal4/UAS-gbb^9.9^; dSod1^G85R^:* 1.25 ± 0.04s, n=19; p=3.9×10^−6^, one-sided t-test with Holm-Bonferroni correction).

Consistent with a previous report showing that hyperactivation of BMP signaling can negatively affect neurons (Nahm et al., 2013), we found that expressing *gbb* using *OK371-Gal4* led to a reduction in wild type larval locomotion (Figure 3B) (*OK371-Gal4/UAS-gbb^9.9^; dSod1^WTLoxP^* = 49.5 ± 1.9mm, n=45; *dSod1^WTLoxP^* = 63.0 ± 2.1mm, n=63; p=1.9×10^−5^, t-test with Holm-Bonferroni correction) as well as a reduction in the number of adults that eclose (Table 1) (*dSod1^WTLoxP^* vs *OK371-Gal4/UAS-gbb^9.9^;dSod1^WTLoxP^* p=0.014). Interestingly, despite the negative impact of overexpressed *gbb* in wild type, we found that *OK371-Gal4* driven *gbb* improved *dSod1^G85R^* larval locomotion and enabled the eclosion of some *dSod1^G85R^* flies (Table 1 and Figure 3A-E).

Bursts of neural activity that correlate with segmental muscle contractions in larvae can be monitored in fictive crawling preparations in which the motor circuit of the dissected larva is left intact (Cattaert et al., 2001 ; Fox et al., 2006). Wandering 3^rd^ instar larvae were fileted, exposing the VNC and its connections through segmentally organized nerves to the body wall musculature. The concordance of muscle contraction and neural activity is evident when bursts of electrical activity are recorded from a single segmental nerve bundle using an extracellular suction electrode. Each burst corresponds to one round of muscle contractions for that body hemisegment as the larval preparation ‘crawls’ *ex vivo*. Consistent with the reduced crawling behavior exhibited by *dSod1^G85R^* larvae, we found that the bursts of neural activity in fictive crawling preparations of *dSod1^G85R^* were less frequent than in controls (Figure 3F and G). When *gbb* expression was driven by *OK371-Gal4* in *dSod1^G85R^*, the bursts of neural activity were more frequent (Figure 3F and G) (*dSod1^WTLoxP^:* 4.0 ± 0.5 events/min, n=18. *dSod1^G85R^*: 1.8 ± 0.3 events/min, n=16. *OK371 -Gal4/UAS-gbb^9.9^; dSod1^G85R^:* 3.5 ± 0.7 events/min, n=14. *dSod1^WTLoxP^* vs *dSod1^G85R^* p=7.3×10^−4^; *dSod1^G85R^* vs *OK371-Gal4/UAS-gbb^9.9^; dSod1^G85R^* p=0.029, unequal variance t-tests with Holm-Bonferroni correction). These fictive crawling measurements paralleled our *in vivo* larval locomotion analyses, demonstrating that the *dSod1^G85R^* crawling defect and its rescue by *OK371-Gal4* driven *gbb* can be observed and recorded *ex vivo*.

### Minor NMJ defects detected in dSod1^G85R^ larvae unlikely to account for reduced locomotion

Despite the abnormal locomotor behavior of *dSod1^G85R^* larvae (Figure 3), we did not find a substantial change in the structure or function of NMJs in mutant 3^rd^ instar larvae (Figure 4). This is in contrast to the severe disruption of NMJ structure and function observed in abdominal muscles of *dSod1^G85R^* pharates (Figure 1). In *dSod1^G85R^* larvae, the number of boutons at muscle 6/7 NMJs is not different from wild type (Figure 4A) (*dSod1^WTLoxP^* = 91.7 ± 2.5 boutons, n=22; *dSod1^G85R^* = 99 ± 3.13 boutons, n=22; p=0.14, t-test with Holm-Bonferroni correction). Similarly, the electrophysiological properties of the *dSod1^G85R^* muscle 6 NMJ did not reveal differences in the amplitude of excitatory events (mEPSPs, eEPSPs, eEPSCS) or the kinetics of evoked excitatory post-synaptic currents (eEPSCs) compared to wild type (Figure 4B-E) <u>(mEPSP amplitude</u>: *dSod1^WTLoxP^* = 0.86 ± 0.09mV, n=12; *dSod1^G85R^* = 0.98 ± 0.05mV, n=18; p=0.23, t-test; <u>eEPSP amplitude</u>: *dSod1^WTLoxP^* = 38.5 ± 2.1mV, n=12; *dSod1^G85R^* was 38.9 ± 1.7mV, n=18; p=0.87, t-test; <u>eEPSC amplitude</u>: *dSod1^WTLoxP^* = −90 ± 7.8nA, n=8; *dSod1^G85R^* = − 91 ± 8.8nA, n=12; p=0.96, t-test; <u>eEPSC rise time</u>: *dSod1^WTLoxP^* = 0.93 ± 0.07ms, n=7; *dSod1^G85R^* = 1.02 ± 0.10ms, n=12; p=0.50, t-test; <u>eEPSC decay time</u>: *dSod1^WTLoxP^* = 9.3 ± 0.64ms, n=8; *dSod1^G85R^* = 9.7 ± 0.60ms, n=12; p=0.65, t-test). We did observe a slight increase in the frequency of mEPSPs in *dSod1^G85R^*, suggesting a mild defect in the *dSod1^G85R^* NMJ (Figure 4B,C) <u>(mEPSP frequency</u>: *dSod1^WTLoxP^* = 2.5 ± 0.38Hz, n=12; *dSod1^G85R^* = 3.6 ± 0.32Hz, n=18; p=0.04, t-test). Interestingly, overexpressing *gbb* in *dSod1^G85R^* motor neurons did not lead to an increase in bouton number, a phenotype typically seen in wild type NMJs when *gbb* expression is driven in motor neurons (James and Broihier, 2011) (Figure 4A) *OK371-Gal4/UA S-gbb^9.9^;dSod 1^WTLoxP^* = 115.0 ± 3.7 boutons, n=15 vs *dSod1^WTLoxP^* = 91.7 ± 2.5 boutons, n=22; p=2.7×10^−5^, t-test with Holm-Bonferroni correction; *OK371-Gal4/UAS-gbb^9.9^;dSod1^G85R^:* 102.3 ± 4.2 boutons, n=13; *dSod1^G85R^:* 99 ± 3.13 boutons, n=22; p=0.26, t-test with Holm-Bonferroni correction), suggesting that *dSod1^G85R^* NMJs are defective in some manner, consistent with their altered mEPSP frequency. While *dSod1^G85R^* larval NMJs may be slightly defective, the fact that the excitatory event amplitudes remain normal suggests that the reduced locomotion of *dSod1^G85R^* larvae is unlikely to result from an NMJ defect.

Since larval muscle contraction requires a coordinated response from the muscle as well as repeated motor neuron firing and sufficient neurotransmitter release (Cattaert et al., 2001), we tested for the possibility that muscle contractility itself is abnormal in *dSod1^G85R^*. We measured the response of muscles 6/7 to a range of motor neuron stimulation frequencies. No alterations in contraction were observed between *dSod1^G85R^* and wild type 3^rd^ instar larvae (Figure 4F and G) (*dSod1^WTLoxP^* n=10 and *dSod1^G85R^* n=10; GLME: genotype effect p= 0.21, stimulus effect p=1.5×10^−28^). Thus, the reduced locomotion of *dSod1^G85R^* 3^rd^ instar larvae is not easily attributed to abnormalities at the NMJ and/or defects in muscle contraction.

### dSod1^G85R^ larval motor neurons exhibit mild hypoexcitability

Alterations in motor neuron excitability have been observed in ALS patients and in some ALS models (Kanai et al., 2006; Iwai et al., 2016; King et al., 2016). Given the relatively minor NMJ defects detected in *dSod1^G85R^* larvae, we investigated the possibility that altered motor neuron excitability could account for the observed reduction in locomotor activity. Patch recordings followed by current steps revealed a slight reduction in the excitability of aCC motor neurons in *dSod1^G85R^* compared to *dSod1^WTLoxP^* (Figure 5A,B) (*dSod1^WTLoxP^* n=13 and *dSod1^G85R^* n=10; GLME: genotype effect p=3.9×10^−6^, stimulus effect p=5.4×10^−80^). While we did not detect differences in resistance, capacitance, voltage threshold, or action potential dynamics between *dSod1^G85R^* and *dSod1^WTLoxP^*, a slightly increased but not statistically significant, current threshold was observed in *dSod1^G85R^* neurons that could account for their lower excitability (Figure 5C-G) (*<u>dSod1^WTLoxP^:</u>* resistance = 0.74 ± 0.08GΩ, capacitance = 27.1 ± 3.0pF, current threshold = 18.72 ± 2.69pA, voltage threshold = −29.3 ± 1.3 mV, n=13; *<u>dSod1^G85R^:</u>* resistance = 0.81 ± 0.08GΩ, capacitance = 22.2 ± 2.8pF, current threshold = 27.58 ± 5.90pA, voltage threshold = − 28.0 ± 2.1 mV, n=10; resistance p=0.57, capacitance p=0.25, current threshold p=0.15, voltage threshold p=0.57, t-tests).

After the current step protocol, aCC motor neurons were voltage clamped at the chloride reversal potential (-70mV) to isolate excitatory post synaptic currents. No major changes in EPSC amplitude or frequency were apparent, indicating that excitatory signals from the central pattern generator (CPG) interneurons must not be different in *dSod1^G85R^* compared to wild type (Figure 5H and I) (*dSod1^WTLoxP^* = 94.2 ± 18.6pA/pF, 11.1 events/min, n=9; *dSod1^G85R^* = 97.8 ± 18.6pA/pF, 11.7 ± 3.3 events/min, n=10; amplitude p=0.84, Wilcoxon rank sum test; frequency p=0.88, unequal variance t-test). In summary, our electrophysiological analysis indicates that while *dSod1^G85R^* aCC motor neurons exhibit slight hypoexcitablity, they receive similar excitatory inputs from CPG interneurons in both *dSod1^WTLoxP^* and *dSod1^G85R^* with an output not distinguishable in terms of eEPSP amplitude or in maximal muscle contraction (Figs. 4 and 5). Hence, no alterations in motor neuron properties were found that could easily account for the reduced motor activity observed in *dSod1^G85R^* larvae.

### Peripheral feedback dysfunction is linked to a reduction in motor output in dSod1^G85R^

The crawling behavior of larvae depends on output from CPG interneurons to generate the rhythmic motor patterns executed by motor neurons and muscles. The rate of the rhythmic signals produced by the CPG can then be modified by feedback from peripheral proprioceptors (Hughes and Thomas, 2007; Song et al., 2007; Berni et al., 2012). Any change in motor circuit output could arise from a disruption in the function of the motor neuron, NMJ, or muscle, or from defects in other circuit components (Hughes and Thomas, 2007; Song et al., 2007). Targeted electrophysiological measurements failed to uncover defects in *dSod1^G85R^* motor neuron and NMJ function (Figs. 4 and 5) that correlate with the clear reduction in neural activity evident in fictive crawling preparations (Figure 3F,G). As such, we considered the possibility that some aspect of the motor pattern initiated by the CPG is abnormal in *dSod1^G85R^*, or that feedback from the periphery to the central nervous system (CNS) is defective. To distinguish between these possibilities, we compared the activity of an intact motor circuit to the activity generated by the CPG when the CNS is separated from, and no longer receives, proprioceptive or other sensory feedback.

Extracellular recordings were obtained from a segmental nerve in intact (I) fictive crawling preparations of *dSod1^WTLoxP^* and *dSod1^G85R^* 3^rd^ instar larvae (Figure 6). To eliminate peripheral feedback, segmental nerves that contain both motor and sensory axonal tracts were cut (C) distally (deafferented) to prevent sensory input to the CNS. Extracellular recordings were then obtained proximally from axonal tracts extending out of the VNC. First, to exclude the possibility that sequential recordings could affect data quality due to a run-down effect, we compared the frequency of activity bursts from the proximal region of segmental nerves in freshly dissected CNS samples to those recorded from CNS samples which had already been recorded from as part of the intact circuit. We found no difference in their activity indicating no evidence of run-down in this experimental paradigm (Figure 6D) (*dSod1^WTLoxP^* F = 4.8 ± 1.2 events/min, n=6; *dSod1^WTLoxP^* C = 3.6 ± 0.4 events/min, n=9; p=0.41, unequal variance t-test).

The frequency of activity bursts detected by extracellular recordings before and after severing the segmental nerve were compared between *dSod1^WTLoxP^* and *dSod1^G85R^* (Figure 6E and F). While a change in event frequency was not observed in *dSod1^WTLoxP^*, the event frequency of *dSod1^G85R^* doubled after deafferenting (Cut-Intact: *dSod1^WTLoxP^* = −0.94 ± 0.59 events/min, n=9, p=0.15, paired t-test; *dSod1^G85R^* = 2.5 ± 0.6 events, n=7, p=0.0054, paired t-test). The doubling of the burst frequency following the removal of ascending input from the periphery, suggested that, in an intact *dSod1^G85R^* animal, sensory feedback slows the motor pattern produced by the CPG. When this apparently defective peripheral feedback is eliminated by severing the segmental nerve, the motor pattern in *dSod1^G85R^* speeds up.

When we compared the rhythmic motor pattern produced by the CPG between *dSod1^WTLoxP^* vs *dSod1^G85R^*, as measured from the isolated CNS, we found a higher rate of bursting in *dSod1^G85R^* compared to *dSod1^WTLoxP^* (Figure 6E cut (C)) (*dSod1^WTLoxP^* C = 3.6 ± 0.4 events/min, n=9; *dSod1^G85R^* C = 5.6 ± 0.7 events/min, n=7, *dSod1^WTLoxP^* Cut vs *dSod1^G85R^* Cut: p=0.041, t-test with Holm-Bonferroni correction) indicating that two components of the *dSod1^G85R^* larval motor circuit exhibit altered functions. Not only does sensory feedback slow motor circuit output but the *dSod1^G85R^* CPG appears to fire more frequently than the control *dSod1^WTLoxP^* CPG. While this higher CPG output would be predicted to increase the frequency of muscle contraction and rate of *dSod1^G85R^* larval locomotion, we observe that when the circuit is intact the defect in peripheral feedback appears to override this CPG increase and slows the overall motor circuit output, resulting in reduced locomotor activity typical of *dSod1^G85R^* larvae.

We have shown that expressing *gbb* in glutamatergic neurons under the control of *OK371-Gal4* can increase *dSod1^G85R^* larval locomotion (Figure 3), and consistent with this suppression of *dSod1^G85R^-*induced motor dysfunction, we found that the event frequency recorded from segmental nerves in the intact preparations of *OK371>gbb; dSod1^G85R^* was higher than that observed in *dSod1^G85R^* (Figure 6B and E intact (I)) (*dSod1^G85R^* I = 3.1 ± 0.4 events/min, n=7; *OK371-Gal4/UAS-gbb^9.9^; dSod1^G85R^* I = 4.4 ± 0.2 events/min, n=6, p=0.014, one-tailed t-test with Holm-Bonferroni correction). Removing peripheral feedback by deafferenting *OK371>gbb; dSod1^G85R^* preparations did not produce a significant difference in the frequency of bursts when compared to the intact preparation (Figure 6E,F), indicating that when *gbb* is expressed in motor neurons and other glutamatergic neurons, sensory feedback is restored (*OK371-Gal4/UAS-gbb^9.9^; dSod1^G85R^* = −1.1 ± 0.5 events, n=6, p=0.064, paired t-test). These findings are consistent with a model in which activation of BMP signaling affects the efficacy of sensory feedback in *dSod1^G85R^* to improve motor function.

### Neuronal processes are altered in the dSod1^G85R^ larval nerve cord

Defects in peripheral feedback could arise from dysfunction in the ascending sensory neurons themselves and/or from a disruption in the processing of sensory input by the interneuron network within the *dSod1^G85R^* VNC. Proprioceptors play a critical role in relaying peripheral feedback, and disrupting normal proprioceptor function slows the larval locomotor pattern by increasing peristaltic wave duration (Hughes and Thomas, 2007; Song et al., 2007). Because our data indicate that peripheral feedback slows *dSod1^G85R^* locomotion and peristaltic wave duration is increased in *dSod1^G85R^*, we considered the possibility that the proprioceptors are defective (Figure 7). *2-21-Gal4* is expressed in proprioceptors (Hughes and Thomas, 2007), whose cell bodies reside on the body wall and project axons into the medial region of the nerve cord where they are thought to synapse with interneurons (Figure 7A) (Merritt and Whitington, 1995; Grueber et al., 2007; Schneider-Mizell et al., 2016). The dendritic arbors of proprioceptors could be visualized in *2-21-Gal4>mRFP* and no gross abnormalities were apparent in *dSod1^G85R^*. The dendritic branching of the ddaE proprioceptor was examined in detail by Sholl analysis in both *dSod1^WTLoxP^* and *dSod1^G85R^* larvae. The ddaE cell body was recognized in the larval body wall based on its position (Figure 7B). The number of ddaE dendritic intersections changed with distance from the cell body, but no significant difference was observed between genotypes, suggesting that in *dSod1^G85R^*, ddaE dendritic projections are not different from *dSod1^WTLoxP^* (Figure 7B; GLME: genotype effect p=0.10, radius length effect p=2.4×10^−13^).

In addition to its expression in proprioceptors, *2-21-Gal4* is also expressed in a subset of interneurons within the nerve cord (Figure 7C; GFP-positive cell bodies in *2-21-Gal4>GFP*). *2-21-Gal4*-positive axonal projections in the longitudinal tracts adjacent to the VNC midline are clearly visualized by mRFP (Figure 7D). The intensity of RFP within the boxed regions of the longitudinal tracts was measured in *dSod1^WTLoxP^* and *dSod1^G85R^* VNCs (Figure 7D). The mRFP signal is less intense in *dSod1^G85R^*suggestive of a reduction in *2-21-Gal4-*labelled neuronal processes (*dSod1^WTLoxP^* = 3.04 ± 0.29AU, n=8; *dSod1^G85R^* = 2.11 ± 0.13AU, n=8, p=0.017, unequal variance t-test). Such a change in *2-21-Gal4* labelled neuronal processes in the *dSod1^G85R^* larval VNC is consistent with our finding that non-motor neuron function is compromised in *dSod1^G85R^* (Figure 6).

### BMP signaling acts in specific components of the motor circuit to rescue dSod1^G85R^-induced dysfunction

Gbb, the BMP7 ortholog has been shown to act as a retrograde signal at the *Drosophila* larval NMJ (McCabe et al., 2003). Transduction of BMP signaling results in phosphorylation of the R-Smad, Mad, in *Drosophila* (Hoodless et al., 1996; Tanimoto et al., 2000). As expected, pMad is detected in motor neuron cell bodies in the VNC, as well as at low levels at the NMJ of wild type (Fuentes-Medel et al., 2012). In *dSod1^G85R^*, the level of pMad in the motor neuron cell body is not different from wild type, but a significant increase in pMad is apparent at the *dSod1^G85R^* NMJ (Figure 8A,B) <u>(MN cell body</u>: *dSod1^WTLoxP^:* 1.0 ± 0.12AU, n=9; *dSod1^G85R^:* 1.2 ± 0.79AU, n=10; p=0.18, t-test. NMJ <u>log2(pMad/HRP) ratio</u>: *dSod1^WTLoxP^* = −3.1 ± 0.24, n=9; *dSod1^G85R^* = −2.1 ± 0.28, n=11; *dSod1^WTLoxP^* vs *dSod1^G85R^* p=0.023, unequal variance t-test with Holm-Bonferroni correction). The consequence of an increase in pMad at the *dSod1^G85R^* NMJ is not yet clear, but this observation, in addition to the fact that an increase in *gbb* does not lead to an overgrowth of *dSod1^G85R^* NMJ (Figure 4A), suggests that the NMJ is not completely normal despite maintaining relatively normal function.

The presence of pMad is an indicator of cells that have received a BMP signal. In response to *gbb* overexpression in glutamatergic neurons (*OK371>gbb; dSod1^G85R^*) we found very high levels of pMad in cells of the VNC outside of the medially located motor neurons (Figure 8A). While *OK371-Gal4* is expressed in glutamatergic motor neurons and premotor interneurons (Figure 8C), the broad distribution of pMad suggests that *OK371>gbb* may result in a non-autonomous activation of BMP signaling due to the secreted nature of Gbb (Figure 8A). Interestingly, despite the expression of *OK371-Gal4* in motor neurons, as clearly indicated by the detection of GFP in the NMJ of *OK371-Gal4>UAS-GFP*, overexpression of *gbb* did not result in a further increase in pMad over the already elevated synaptic pMad in the *dSod1^G85R^* NMJ (Figure 8B) <u>(log_2_(pMad/HRP) ratio</u>: *dSod1^G85R^* = −2.1 ± 0.28, n=11; *OK371-Gal4/UAS-gbb^9.9^; dSod1^G85R^* = −1.8 ± 0.30, n=9. *dSod1^G85R^* vs *OK371-Gal4/UAS-gbb^9.9^; dSod1^G85R^* p=0.50, equal variance t-test with Holm-Bonferroni correction). Thus, it appears that Gbb produced by glutamatergic neurons can induce signaling in a non-autonomous manner within the VNC, but it is unable to induce a change in pMad levels at the motor neuron synapse. This inability of Gbb to induce synaptic pMad is in agreement with the findings of Sulkowski et al. (2016). However, the ability of Gbb to induce high levels of pMad in a non-autonomous manner within cells of the VNC raises the possibility that Gbb contributes to the rescue of *dSod1^G85R^* dysfunction by activation of BMP signaling in other cell types, such as non-motor neurons.

To better define in which cells activation of BMP signaling is critical for the suppression of *dSod1^G85R^*-associated phenotypes, we induced the pathway in a cell autonomous manner by expressing a constitutively active form of the *Drosophila* BMP type I receptor, SaxA (Xie et al., 1994; Twombly et al., 2009; Ball et al., 2010; Piccioli and Littleton, 2014)(Figure 9A,B). Activation of BMP signaling in either glutamatergic neurons (*OK371-Gal4>SaxA; dSod1^G85R^*), which include motor neurons and a subset of premotor interneurons, or in cholinergic neurons, which make up sensory and interneurons in *Drosophila* (*ChAT-Gal4>SaxA; dSod1^G85R^*), leads to an increase in the locomotor activity of *dSod1^G85R^* larvae (Figure 9A,B) (*dSod1^WTLoxP^* = 48.9 ± 3.2mm, n=24; *dSod1^G85R^* = 19.9 ± 1.9mm, n=22; *OK371-Gal4/+;UAS-SaxA, dSod 1 ^G85R^/dSod 1^G85R^* = 45.2 ± 3.2mm, n=40; *OK371-Gal4/+;dSod1^G85R^* = 29.7 ± 1.8mm, n=42; *UAS-SaxA,dSod1^G85R^/dSod1^G85R^* = 29.1 ± 3.7mm, n=19; *dSod1^WTLoxP^* vs *dSod1^G85R^* p=1.3×10^−8^; *OK371-Gal4/+;UAS-SaxA,dSod1^G85R^/dSod1^G85R^* vs *dSod1^G85R^* p=1.7×10^−8^; *OK371-Gal4/+;UAS-SaxA,dSod1^G85R^/dSod1^G85R^* vs *OK371-Gal4/+;dSod1^G85R^* p=1.4×10^−4^; *OK371-Gal4/+;UAS-SaxA,dSod1^G85R^/dSod1^G85R^* vs *UAS-SaxA,dSod1^G85R^/dSod1^G85R^* p=3.3×10^−3^, t-tests with Holm-Bonferroni corrections. *dSod1^WTLoxP^* = 57.3 ± 3.6mm, n=25; *dSod1^G85R^* = 30.8 ± 3.3mm, n=22; *ChAT-Gal4/+;UAS-SaxA,dSod1^G85R^/dSod1^G85R^* = 49.7 ± 3.3mm, n=42; *ChAT-Gal4/+;dSod1^G85R^* = 36.4 ± 3.3mm, n=32; *UAS-SaxA,dSod1^G85R^/dSod1^G85R^* = 32.3 ± 3.9mm, n=19; *dSod1^WTLoxP^* vs *dSod1^G85R^* p=9.7×10^−6^; *ChAT-Gal4/+;UAS-SaxA,dSod1^G85R^/dSod1^G85R^* vs *dSod1^G85R^* p=1.6×10^−3^; *ChAT-Gal4/+;UAS-SaxA,dSod1^G85R^/dSod1^G85R^* vs *ChAT-Gal4/+;dSod 1^G85R^* p=6.7×10^−3^; *ChAT-Gal4/+;UAS-SaxA,dSod1^G85R^/dSod1^G85R^* vs *UAS-SaxA,dSod1^G85R^/dSod1^G85R^* p=5.6×10^−3^; t-tests with Holm-Bonferroni corrections).

Activation of BMP signaling in cholinergic neurons also resulted in an increase in successful eclosion, with 10.2% of *dSod1^G85R^* adults (*ChAT-Gal4/+; UAS-SaxA dSod^G85R^/dSod^G85R^*) emerging from their pupal cases (Table 2) (*dSod1^WTLoxP^* vs *dSod1^G85R^* p=3.1*×*10^−101^; *ChAT-Gal4/+;UAS-SaxA,dSod1^G85R^/dSod1^G85R^* vs *dSod1^G85R^* p=1.5*×*10^−4^; *ChAT-Gal4/+;UAS-SaxA,dSod1^G85R^/dSod1^G85R^* vs *ChAT-Gal4/+;dSod1^G85R^* p=8.6*×*10^−6^; *ChAT-Gal4/+;UAS-SaxA,dSod1^G85R^/dSod1^G85R^* vs *UAS-SaxA,dSod1^G85R^/dSod1^G85R^* p=1.8*×*10^−3^; Fisher’s Exact tests with Holm-Bonferroni corrections).

Expression of *SaxA* in multidendritic sensory neurons (*MD-Gal4*) did not rescue either *dSod1^G85R^* larval locomotion or eclosion (Figure 9 and Table 2) (Larval locomotion: *dSod1^WTLoxP^* = 48.2 ± 1.8mm, n=60; *dSod1^G85R^* = 27.8 ± 2.1mm, n=45; *MD-Gal4/+;UAS-SaxA, dSod1^G85R^/dSod1^G85R^* = 33.3 ± 2.6mm, n=34; *MD-Gal4/+;dSod1^G85R^* = 29.7 ± 2.4mm, n=35; *UAS-SaxA,dSod1^G85R^/dSod1^G85R^* = 34.0 ± 3.0mm, n=42; *dSod1^WTLoxP^* vs *dSod1^G85R^* p=1.3×10^−10^; *MD-Gal4/+;UAS-SaxA,dSod1^G85R^/dSod1^G85R^* vs *dSod1^G85R^* p=0.32; *MD-Gal4/+;UAS-SaxA,dSod1^G85R^/dSod1^G85R^* vs *MD-Gal4/+;dSod1^G85R^* p=0.65; *MD-Gal4/+;UAS-SaxA,dSod1 ^G85R^/dSod1^G85R^* vs *UAS-SaxA,dSod1^G85R^/dSod1^G85R^* p=0.87; t-tests with Holm-Bonferroni corrections. Eclosion: *dSod1^WTLoxP^* vs *dSod1^G85R^* p=1.3×10^−157^; *MD-Gal4/+;UAS-SaxA,dSod1 ^G85R^/dSod1^G85R^* vs *dSod1^G85R^* p=0.58; *MD-Gal4/+;UAS-SaxA,dSod1^G85R^/dSod1^G85R^* vs *MD-Gal4/+;dSod1^G85R^* p=1.0; *MD-Gal4/+;UAS-SaxA,dSod1^G85R^/dSod1^G85R^* vs *UAS-SaxA,dSod1^G85R^/dSod1^G85R^* p=1.0; Fisher’s Exact tests with Holm-Bonferroni corrections). However, interestingly, driving SaxA in a subset of multidendritic sensory neurons, the proprioceptors (*2-21-Gal4>SaxA; dSod^G85R^*), enabled both an increase in *dSod^G85R^* larval locomotion and adult eclosion (35.6% of *dSod1^G85R^* animals to eclose) (Figure 9B and Table 2) (Larval locomotion: *dSod1^WTLoxP^* = 50.5 ± 3.7mm, n=19; *dSod1^G85R^* = 30.6 ± 4.4mm, n=35; *2-21-Gal4,dSod1^G85R^/UAS-SaxA,dSod1^G85R^* = 48.6 ± 5.1mm, n=30; *2-21-Gal4,dSod1^G85R^/dSod1^G85R^* = 30.8 ± 3.6mm, n=39; *UAS-SaxA,dSod1^G85R^/dSod1^G85R^* = 27.7 ± 3.9mm, n=39; *dSod1^WTLoxP^* vs *dSod1^G85R^* p=2.4×10^−3^; *2-21-Gal4,dSod1^G85R^/UAS-SaxA,dSod1^G85R^* vs *dSod1^G85R^* p=4.8×10^−3^; *2*-*21-Gal4,dSod1^G85R^/UAS-SaxA,dSod1^G85R^* vs *2-21-Gal4,dSod1^G85R^/dSod1^G85R^* p=4.2×10^−3^; *2-21-Gal4,dSod1^G85R^/UAS-SaxA,dSod1^G85R^* vs *UAS-SaxA,dSod1^G85R^/dSod1^G85R^* p=2.4×10^−3^; t-tests with Holm-Bonferroni corrections. Eclosion: *dSod1^WTLoxP^* vs *dSod1^G85R^* p=3.3×10^−218^; *2-21-Gal4,dSod1^G85R^/UAS-SaxA,dSod1^G85R^* vs *dSod1^G85R^* p=1.8×10^−35^; *2-21-Gal4,dSod1^G85R^/UAS-SaxA,dSod1^G85R^* vs *2-21-Gal4,dSod1^G85R^/dSod1^G85R^* p=2.3×10^−30^; *2-21-Gal4,dSod1^G85R^/UAS-SaxA,dSod1^G85R^* vs *UAS-SaxA,dSod1^G85R^/dSod1^G85R^* p=8.6×10^−37^; Fisher’s Exact tests with Holm-Bonferroni corrections). Driving expression of *SaxA* in either muscles (*BG57-Gal4*) or glial cells (*Repo-Gal4*) did not result in a rescue *dSod1^G85R^* larval locomotion or eclosion, indicating that the ability of BMP signaling activation to rescue *dSod1^G85R^* phenotypes in successful in only some cell types and/or with only some Gal4 drivers (Figure 9B and Table 2) (*<u>BG57-Gal4</u>* <u>larval locomotion</u>: *dSod1^WTLoxP^* = 64.6 ± 2.7mm, n=45; *dSod1^G85R^* = 27.2 ± 2.7mm, n=45; *BG57-Gal4, dSod1^G85R^/UAS-SaxA,dSod1^G85R^* = 29.3 ± 2.9mm, n=45; *BG57-Gal4,dSod1^G85R^/dSod1^G85R^* = 26.5 ± 2.8mm, n=38; *UAS-SaxA,dSod1^G85R^/dSod1^G85R^* = 25.7 ± 2.8mm, n=40; *dSod1^WTLoxP^* vs *dSod1^G85R^* p=4.2×10^−15^; *BG57-Gal4,dSod1^G85R^/UAS-SaxA,dSod1^G85R^* vs *dSod1^G85R^* p=1.0; *BG57-Gal4,dSod 1^G85R^/UAS-SaxA,dSod1^G85R^* vs *BG57-Gal4,dSod1^G85R^/dSod1^G85R^* p=1.0; *BG57-Gal4,dSod 1^G85R^/UAS-SaxA,dSod1^G85R^* vs *UAS-SaxA,dSod1^G85R^/dSod1^G85R^* p=1.0; t-tests with Holm-Bonferroni corrections. *<u>BG57-Gal4</u>* <u>eclosion</u> : *dSod1^WTLoxP^* vs *dSod1^G85R^* p<10^−250^; *BG57-Gal4,dSod1^G85R^/UAS-SaxA,dSod1^G85R^* vs *dSod1^G85R^* p=1.0; *BG57-Gal4,dSod1^G85R^/UAS-SaxA,dSod1^G85R^* vs *BG57-Gal4,dSod1^G85R^/dSod1^G85R^* p=1.0; *BG57-Gal4,dSod1^G85R^/UAS-SaxA,dSod1^G85R^* vs *UAS-SaxA,dSod1^G85R^/dSod1^G85R^* p=1.0; Fisher’s Exact tests with Holm-Bonferroni corrections. *<u>Repo-Gal4</u>* <u>larval locomotion</u>: *dSod1^WTLoxP^* = 64.2 ± 2.3mm, n=51; *dSod1^G85R^* = 41.7 ± 2.9mm, n=44; *Repo-Gal4,dSod1^G85R^/UAS-SaxA,dSod1^G85R^* = 36.2 ± 3.6mm, n=33; *Repo-Gal4,dSod1^G85R^/dSod1^G85R^* = 48.2 ± 3.8mm, n=49; *UAS-SaxA,dSod1^G85R^/dSod1^G85R^* = 39.8 ± 3.8mm, n=40; *dSod1^WTLoxP^* vs *dSod1^G85R^* p=9.3×10^−8^; *Repo-Gal4,dSod1^G85R^/UAS-SaxA,dSod1^G85R^* vs *dSod1^G85R^* p=0.48; *Repo-Gal4,dSod1^G85R^/UAS-SaxA,dSod1^G85R^* vs *Repo-Gal4,dSod1^G85R^/dSod1^G85R^* p=0.10; *Repo-Gal4,dSod1^G85R^/UAS-SaxA,dSod1^G85R^* vs *UAS-SaxA,dSod1^G85R^/dSod1^G85R^* p=0.51; t-tests with Holm-Bonferroni corrections. *<u>Repo-Gal4</u>* <u>eclosion</u>: *dSod1^WTLoxP^* vs *dSod1^G85R^* p<10^−250^; *Repo-Gal4,dSod1^G85R^/UAS-SaxA,dSod1^G85R^* vs *dSod1^G85R^* p=3.2×10^−4^; *Repo-Gal4,dSod1^G85R^/UAS-SaxA,dSod1^G85R^* vs *Repo-Gal4,dSod1^G85R^/dSod1^G85R^* p=0.45; *Repo-Gal4,dSod1^G85R^/UAS-SaxA,dSod1^G85R^* vs *UAS-SaxA,dSod1^G85R^/dSod1^G85R^* p=7.4×10^−3^; Fisher’s Exact tests with Holm-Bonferroni corrections).

Taken together, motor defects associated with *dSod1^G85R^* can be rescued by the cell-autonomous activation of BMP signaling in at least two different components of the motor circuit, glutamatergic and cholinergic neurons. Within the cholinergic population, we found robust suppression of *dSod1^G85R^* motor defects by the specific activation of BMP signaling in proprioceptors (*2-21-Gal4*), emphasizes the importance of this class of neurons in the restoration of motor circuit function. Importantly, this result, together with our findings of a defect in sensory feedback that underlies the early circuit dysfunction of *dSod1^G85R^*, underscores the importance of non-motor neurons in the progression of ALS-associated neurodegeneration. Furthermore, our study highlights sensory neurons as a source for the identification of biomarkers indicative of the early disease state and as a site for the development of therapeutics.

## Discussion

A hallmark of late-stage ALS is the profound degeneration of motor neurons, however, the site and initiating events responsible for the ultimate degeneration of motor neurons per se are not fully understood. Significant progress identifying cellular defects associated with fALS has been made, as have attempts to reduce degeneration (Boillée et al., 2006; Yamanaka et al., 2008; Casci and Bhan, 2015; Freibaum et al., 2015; Jovičić et al., 2015; Zhang et al., 2015; Nagy et al., 2016). In mice, motor neuron death can be prevented by blocking motor neuron expression of mutant FUS or TDP43 (Ditsworth et al., 2017; Scekic-Zahirovic et al., 2017). Curiously, while motor neuron death was prevented, locomotion was still impaired, raising the possibility that cells other than motor neurons can affect the onset and/or progression of ALS (Pramatarova et al., 2001; Boillée et al., 2006; Jaarsma et al., 2008; Yamanaka et al., 2008). Glial cells have also been shown to influence ALS progression (Di Giorgio et al., 2007; Lobsiger and Cleveland, 2007). Here, using a knock-in SOD1-ALS model, we report that non-motor neuron dysfunction in early stage animals correlates with disrupted locomotion in the absence of motor neuron degeneration (Figure 10).

**Figure 10.**
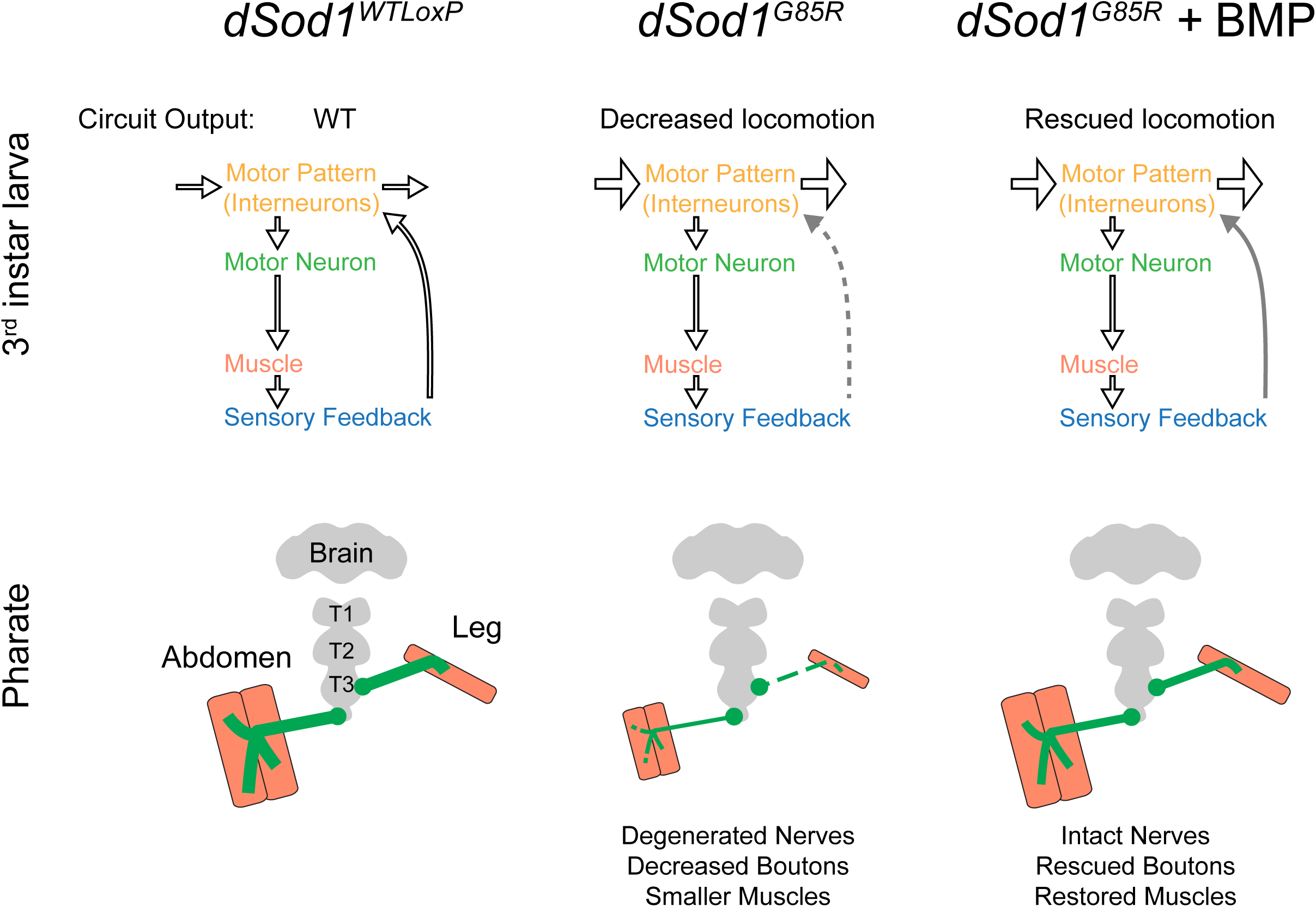
Graphical summary. Defects in *dSod1^G85R^* motor circuitry are rescued by activation of BMP signaling. **Top:** Schematic representation of larval motor circuit. The larval locomotor pattern (yellow) is dictated by both CPG activity (horizontal arrows) in the ventral nerve cord (VNC), signals from sensory neurons (blue) in the periphery and their integration by interneurons (arrow to motor pattern) to provide feedback to the CPG. Early stage *dSod1^G85R^* animals (3^rd^ instar larvae) exhibit reduced locomotion but no degeneration of significant defects in motor neuron (green) function. Instead, an increase in CPG activity (wide horizontal arrows) in *dSod1^G85R^* was revealed along with a disruption to peripheral feedback (dotted gray line) that results in reduced locomotor activity. Activation of BMP signaling increases *dSod1^G85R^* larval locomotion. The heightened activity of *dSod1^G85R^* CPG is unchanged but feedback from the periphery is improved (solid gray arrow). Cell autonomous activation of BMP signaling in either glutamatergic neurons, cholinergic neurons, or proprioceptors increases *dSod1^G85R^* locomotion. **Bottom:** Adult *Drosophila* abdominal muscles and leg muscles are innervated by different sets of motor neurons (green). Both sets of motor neurons have disrupted morphology in end stage *dSod1^G85R^* animals. Synaptic output at the abdominal NMJ, as well as muscle size are decreased. These abnormalities in motor neuron morphology, motor neuron output, and muscle size likely contribute to *dSod1^G85R^* eclosion failure. The expression of *gbb* in *dSod1^G85R^* motor neurons improves motor neuron morphology, motor neuron output, muscle size, and allows the eclosion of some mutant animals. Driving expression of *SaxA* in cholinergic neurons or proprioceptors increases *dSod1^G85R^* larval locomotion and the percentage of animals that eclose.

End-stage *dSod1^G85R^* knock-in animals exhibit key features of ALS, including severely compromised locomotion, reduced motor neuron function, visible motor neuron degeneration, and decreased muscle size (Şahin et al., 2017; Figure 10). Earlier, at the first sign of compromised locomotor activity, we found the functional properties of *dSod1^G85R^* motor neurons to be largely normal. Circuit analysis revealed a specific disruption in the function of non-motor neurons, implicating this dysfunction as an indicator of early ALS. We also found that both early and late stage *dSod1^G85R^* motor defects can be attenuated by BMP signaling, a pathway known to stimulate neuronal growth in both invertebrates and vertebrates (Lee-Hoeflich et al., 2004; Sun et al., 2007; Hocking et al., 2009). Given the disruption in sensory feedback in the *dSod1^G85R^* larval motor circuit, it is noteworthy that both compromised locomotor activity of mutant larvae and eclosion failure of mutant adults is suppressed by activating BMP signaling specifically within proprioceptors, the subset of sensory neurons that relay the status of muscle contraction back to the CNS (Figure 10). These findings, coupled with those that have previously implicated non-autonomous factors in ALS degeneration (Ilieva et al., 2009), strongly support a hypothesis that non-motor neurons, such as proprioceptors, are impacted first in ALS and their dysfunction ultimately leads to motor neuron degeneration

### Motor circuit dysfunction results in locomotor defects in a knock-in model of Sod1-ALS

Although compromised non-motor neurons have not been formally shown to cause motor function loss in ALS, the suggestion that neurodegenerative disorders can arise from circuit or network disorders has been made (Warren et al., 2013; Fornito et al., 2015). Certainly, in late stage ALS, disruptions in interneuron and/or sensory function were found to accompany severe motor deficits (Theys et al., 1999; Fischer et al., 2005; Stephens et al., 2006; Isaacs et al., 2007; Pugdahl et al., 2007; Sunico et al., 2011; McGown et al., 2013; Vinsant et al., 2013; Vaughan et al., 2015; Clark et al., 2017). Interestingly, in several models of ALS an observed locomotor defect is associated with only a minor decrease in NMJ transmission (Diaper et al., 2013; Rocha et al., 2013; Zhang et al., 2015), and one that would not have been expected to overcome the large safety factor observed at NMJs (Marrus and DiAntonio, 2005).

Recent studies on the motor neuron disease Spinal Muscular Atrophy (SMA), highlighted sensory and interneurons in disease progression. Interneurons deficient in the *SMN1* gene were found to increase the resistance and excitability of co-cultured wild type motor neurons in a stem-cell based SMA model (Simon et al., 2016). In *Drosophila*, the *smn* gene is required in both proprioceptive sensory neurons and interneurons for proper motor neuron function (Imlach et al., 2012; Lotti et al., 2012). In mice, expression of SMN in proprioceptors was shown to dramatically influence motor neuron excitability (Fletcher et al., 2017). The disruption in the knock-in *dSod1^G85R^* ALS model of sensory and interneuron function resembles that found in models of SMA. Future studies to assess whether motor neuron and muscle deficits in ALS arise from a specific loss of proprioceptive function will be of interest. Certainly, our findings presented here, coupled with those in SMA models, support the narrative that circuitry disruption is an important consideration in motor neuron disease.

### Activation of BMP signaling suppresses ALS phenotypes

In *Drosophila*, BMP signaling contributes to NMJ growth, neurotransmission, and synaptic plasticity (McCabe et al., 2003; Baines, 2004; Goold and Davis, 2007; Berke et al., 2013; Piccioli and Littleton, 2014). An increase in BMP signaling is sufficient to induce NMJ growth and produce more robust eEPSPs (James and Broihier, 2011; James et al., 2014; our observations). Increased BMP signaling due to a reduction in gene dosage of the Dad antagonist was shown to alleviate some NMJ abnormalities in a *Drosophila* SMA model, and more recently in an ALS model (Chang et al., 2008; Deshpande et al., 2016). We initially explored the possibility that activating BMP signaling could ameliorate motor neuron degeneration in *dSod1^G85R^* pharates. Indeed, driving the expression of the BMP ligand, Gbb, in motor neurons and glutamatergic interneurons rescued adult *dSod1^G85R^* phenotypes at the NMJ. But even in early-stage *dSod1^G85R^* animals where NMJ defects are minimal, we found that *gbb* expression in glutamatergic neurons could still restore locomotion, presumably by acting somewhere other than the NMJ. Phenotypic suppression of *dSod1^G85R^* was not only achieved by cell-autonomous activation of BMP signaling glutamatergic motor neurons and interneurons, but also in cholinergic sensory and interneurons. The finding that non-motor neuronal processes appear reduced in *dSod1^G85R^* points to these cells as a potential source of dysfunction in the ALS motor circuit. Attempts to record directly from the proprioceptors were unsuccessful but the rescue of *dSod1^G85R^* by SaxA expression in a subset of sensory neurons, the proprioceptors, is concordant with the results of our circuit analysis highlighting a disruption in sensory feedback as the earliest defect detected in *dSod1^G85R^*, and supports the hypothesis that non-motor neurons play a critical role in ALS motor dysfunction. Curiously, expressing *SaxA* in all multidendritic sensory neurons did not rescue while both broader expression of *SaxA* in the cholinergic neurons (sensory and interneurons), and specific expression in the proprioceptive sensory neurons, resulted in restoration of function. This could reflect a differential response to BMP signaling (ie. Follansbee et al., 2017). Thus, in the case of broad activation in all cholinergic neurons the cumulative effect may on balance be positive to result in an increase in locomotion and eclosion.

Interestingly, several pieces of evidence indicate that BMP signaling itself is disrupted in *dSod1^G85R^*. Normally, the expression of *gbb* in motor neurons results in NMJ growth, a consequence not seen in *dSod1^G85R^* despite the clear indication of active signaling in cells of the VNC showing high levels of pMad. Furthermore, the NMJs of *dSod1^G85R^* show a significant elevation in synaptic pMad compared to controls. Dysregulation of BMP signaling may be a common feature of ALS, as the loss of *Drosophila* TDP43 ortholog, *tbph*, function and overexpression of hTDP43, both associated with NMJ disruptions, exhibit decreased synaptic pMad levels (Deshpande et al., 2016). While disrupted in both SOD1 and TDP43 ALS models, the mechanics of how BMP signaling is regulated likely differs between these ALS models, as manipulation of TDP43 levels results not in an elevation of synaptic pMad but a reduction. Certainly, the molecular mechanisms controlling production of synaptic pMad versus nuclear pMad in the motor neuron cell body appear distinct, as nuclear pMad levels remain unchanged in both the *dSod1^G85R^* and TDP43 models. It is possible that multiple aspects of BMP signaling are defective in ALS. The ligand-independent nature of synaptic pMad production, its uncoupling from NMJ growth, and its proposed role in stabilizing postsynaptic Type IIA glutamate receptors (Sulkowski et al., 2016) raises future research questions of exactly how endogenous BMP signaling is altered in ALS and how selective cellular activation of the pathway leads to restoration of motor function.

Overall, our findings reinforce the importance of developing disease models that recapitulate the endogenous spatial and temporal expression of disease alleles, especially as a complementary approach to targeted expression of orthologous genes. The ability to study a fully ‘diseased’ motor circuit and to probe discrete elements in the context of the intact circuit, is crucial for elucidating the interrelatedness of a system more analogous to the patient. While additional studies are necessary, the work presented here provides a framework that highlights the consequences of disease-associated dysfunction in a relatively simple motor circuit. Importantly, our finding that ALS-like phenotypes can be alleviated by activation of the neurotrophic BMP pathway in non-motor neurons lays the groundwork to identify targets of BMP signaling responsible for such rescue. In the future, developing strategies that target non-motor neurons may provide additional benefits to therapeutic approaches that have focused on motor neurons in the treatment of ALS.

## Acknowledgements

We thank Arturo Andrade and Nara Muraro for electrophysiology training and advice. We are grateful to the Bloomington Drosophila Stock Center (NIH P40OD018537), Michael O’Connor, John Thomas, and Heather Broihier for *Drosophila* lines and sharing reagents. We acknowledge the Developmental Studies Hybridoma Bank (DSHB), created by the NICHD of the NIH and maintained at The University of Iowa, Department of Biology, for antibodies used in this study. We also thank the Leduc Bioimaging facility at Brown University for training, and Lipscombe and Wharton lab members for on-going discussions.

